# Transmembrane exchange of fluorosugars: characterization of red cell GLUT1 kinetics using ^19^F NMR

**DOI:** 10.1101/331231

**Authors:** D Shishmarev, CQ Fontenelle, I Kuprov, B Linclau, PW Kuchel

**Keywords:** fluorinated sugar, glucose transport, NMR spectroscopy, erythrocytes, Michaelis-Menten kinetics, enzyme-substrate interactions

## Abstract

We developed a novel approach for quantifying the equilibrium-exchange kinetics of carrier-mediated transmembrane transport of fluorinated substrates. Our method is based on an adapted kinetic theory describing the concentration dependence of the transmembrane-exchange rates of two simultaneously transported species. Using the new approach, we quantified the kinetics of membrane transport of both anomers of three monofluorinated glucose analogues in human erythrocytes (red blood cells: RBCs) using ^19^F nuclear magnetic resonance (NMR) exchange spectroscopy (EXSY). An inosine-based glucose-free medium was shown to promote survival and stable metabolism of RBCs over the duration of the experiments (a few hours). Earlier NMR studies only yielded the apparent rate constants and transmembrane fluxes of the anomeric species, whereas we were able to categorize the two anomers in terms of the catalytic activity (specificity constants) of the glucose transport protein GLUT1 towards them. Differences in the membrane permeability of the three glucose analogues were qualitatively interpreted in terms of local perturbations in the bonding of substrates to amino-acid residues in the active site of GLUT1. The methodology of this work will be applicable to studies of other carrier-mediated membrane transport processes, especially those with competition between simultaneously transported species. The GLUT1-specific results will apply to the design of probes of glucose transport, or inhibitors of glucose metabolism in cells including those exhibiting the Warburg effect.

**ABBREVIATIONS:**

EXSY: exchange spectroscopy
FDG: fluoro-deoxy-glucose
FDG-*n*: *n*-fluoro-*n*- deoxy-D-glucose (*n* = 2, 3, 4)
FID: free induction decay
Glc: D-glucose
NMR: nuclear magnetic resonance
RBC: red blood cell

## INTRODUCTION

### Glucose transport via GLUT proteins

D-Glucose (Glc) metabolism is essential to all living systems (1). Due to the marked hydrophilicity of Glc, lipid-bilayer membranes impose a significant barrier to its exchange into and out of cells. Consequently, most organisms have evolved specialized membrane proteins that catalyze this process (2). In humans, the GLUT family of integral membrane proteins, consisting of 14 known isoforms (3, 4), perform this function. Of these, GLUT1 (UniProt Accession ID: P11166) is the most abundant and, of relevance to the present work, is the major glucose transporter in the human erythrocyte (red blood cell; RBC), where it is essential for cell viability (5).

GLUT1 and some other GLUT isoforms are overexpressed in cancer cells, in response to meeting the increased demand for Glc that defines the Warburg effect (6, 7). This makes members of this class of proteins appealing as therapeutic drug targets (8, 9), and there is a considerable interest in the development of novel compounds that bind to and inhibit GLUT1 (10, 11). In order to improve the molecular recognition of drug candidates by GLUT proteins, various means of incorporating Glc into larger molecules, or mimicking the Glc scaffold, are being actively explored (12).

The crystal structure of GLUT1 was reported by Deng et al. in 2014 (13). The crystallized protein had *n*-nonyl-β-D-glucopyranoside in the binding pocket, which allowed the transporter to be captured in the inward-open conformation. In 2015, the same group reported a high-resolution structure of GLUT3 complexed with Glc in an outward-occluded conformation (14). Two additional crystal structures of GLUT3 with the exofacial inhibitor maltose, captured in outward-open and outward-occluded conformations, were also reported (14). These developments enabled a schematic interpretation of the molecular recognition forces that govern binding and transport of Glc by the GLUTs. On the other hand, there are few quantitative results that describe these proteins “in action” as they bind and discharge their substrates, thus mediating transmembrane Glc flux.

### F-substituted glucoses as substrates

A fluorine atom can serve as a hydrogen-bond acceptor, but unlike an -OH group, it cannot be a hydrogen-bond donor (15). Therefore, systematic substitutions of -OH groups by H or F atoms in carbohydrates can be used to probe the participation of specific -OH groups in the binding of carbohydrates to enzymes (16-18). Several fluoro-deoxy-glucoses (FDGs) have been assessed as GLUT1 substrates using ^19^F-NMR spectroscopy (19-22). Nevertheless, only apparent rate constants for transport using a single substrate concentration were estimated, without taking into account Michaelis-Menten kinetics and substrate saturation effects. In the present work, we adapted a previously developed kinetic theory and studied the transport of monofluorinated glucoses in human RBCs at a range of different substrate concentrations. This allowed interpretation of the results in terms of the specificity constants of GLUT1 towards these fluorosugars as it operates *in situ*, in the membranes of fully viable RBCs.

## MATERIALS AND METHODS

### Materials

The three studied monofluoro-deoxy-D-glucoses (Fig. 1) were obtained from Carbosynth (Compton, West Berkshire, UK). Stock solutions were prepared in deuterated saline (154 mM NaCl in D_2_O). The 99% pure D_2_O was from the Australian Institute of Nuclear Science and Engineering (Lucas Heights, NSW, Australia).

**FIGURE 1.**
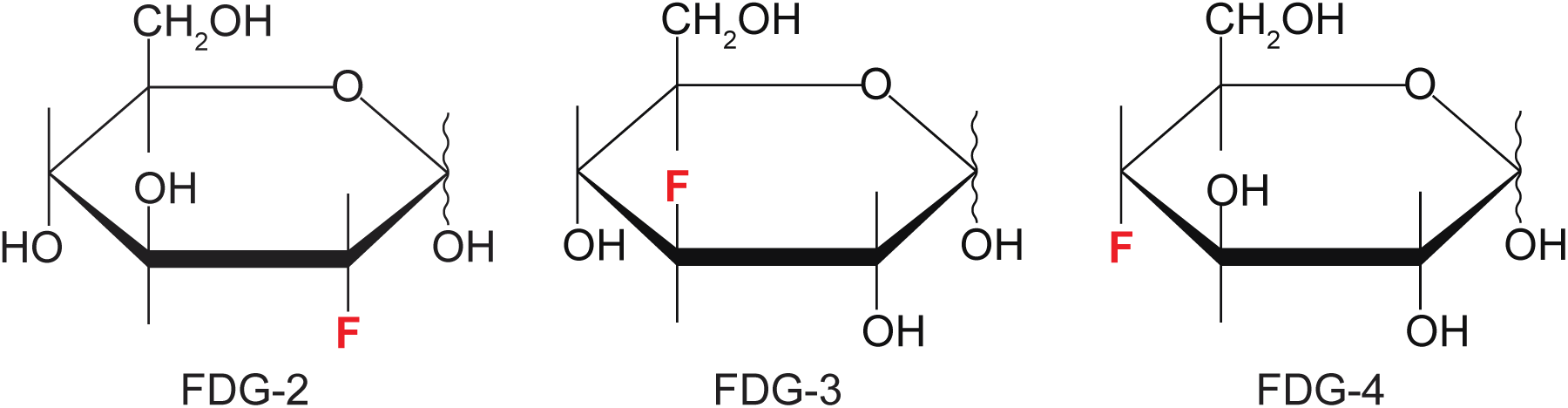
The three studied monofluorinated glucose analogues.

The ‘inosine saline’ medium used for RBC suspensions (300 mOsm kg^-1^; pH 7.4), consisted of 10 mM inosine; 10 mM sodium pyruvate; 5 mM NaH_2_PO_4_; 4.5 mM NaOH; 10 mM KCl and 133 mM NaCl in Milli-Q H_2_O. The osmolality of the saline solutions was measured with a vapor-pressure osmometer (model 5520, Wescor Instruments, Logan, UT, USA). All stock saline solutions were isotonic with RBCs; they were passed three times through a cellulose acetate filter with a pore size of 0.2 µm (Millipore, MA, USA) and stored at 4°C.

Cytochalasin B (cytoB) was purchased from Sapphire Bioscience (Redfern, NSW, Australia). Its 20 mM stock solution was prepared in DMSO-d6 and typically 2.5 µL of this solution was added to 0.5 mL of an NMR sample in GLUT1 inhibition studies; this led to a final concentration of 0.1 mM. All other fine chemicals were from Sigma-Aldrich (St Louis, MO, USA).

### RBC sample preparation

Fresh blood was taken from a single donor by cubital-fossa venipuncture (informed consent was obtained from the subject and the approval given by The University of Sydney Human Ethics Committee). Heparin was used as an anti-coagulant at a concentration of 15 IU (mL blood)^-1^ (23). The blood sample (∼20 mL) was centrifuged at 3000 *g* for 5 min at 10°C, which allowed the separation of RBCs by vacuum-pump aspiration of the blood plasma and buffy coat.

The RBCs were washed twice in saline (154 mM NaCl in Milli-Q H_2_O) and then twice in ‘inosine saline’. Washing involved re-suspending the cells in ∼5 volumes of the washing medium, centrifuging at 3000 *g* for 5 min at 10°C and removing the supernatant by vacuum-pump aspiration. Before the last washing step with saline, the cells were bubbled with carbon monoxide for 10 min to convert the RBC hemoglobin into its stable diamagnetic form for optimal resolution and signal-to-noise in the subsequent NMR spectra.

The obtained RBC suspensions were mixed with various amounts of fluorosugars in 5-mm NMR tubes. In all ^19^F-NMR experiments, the final volume of the sample in an NMR tube (*V*_tot_) was 0.5 mL, which had 10% (v/v) D_2_O and hematocrit (*Ht*) = 60-70%. In each individual case, the *Ht* value was measured using a capillary centrifuge (Clements, North Ryde, NSW, Australia) and taken into account when calculating the total aqueous volume of the sample (Eq. S1 in the Supporting Material). All molar substrate concentrations in the NMR samples were calculated with respect to the total aqueous volume of the sample (*V*_aq_), rather than the total volume of the RBC suspension in the NMR tube (*V*_tot_).

### NMR spectroscopy

^19^F-NMR spectra were acquired at 376.46 MHz on a Bruker Avance III 400 spectrometer (Bruker BioSpin, Karlsruhe, Germany) using either 5-mm dual ^19^F/^1^H or 5-mm BBFO probes (Bruker). ^1^H-NMR spectra were acquired using a 10-mm BBOG probe (Bruker). NMR spectra were recorded and processed using Bruker TopSpin 3.5 software. In all experiments, the sample temperature was 37°C, which was calibrated using a standard sample of neat methanol-d4 (TopSpin command ‘calctemp’).

Figure 2 shows the 1D and 2D exchange spectroscopy (EXSY) pulse sequences that were used. ^1^H decoupling during the indirect and direct acquisition periods was achieved with WALTZ-16 composite-pulse decoupling (24) (90° decoupler-pulse duration of 90 µs centered at 4 ppm), and a high-power 180° ^1^H pulse, respectively. The longitudinal relaxation time constants (*T*_1_) were estimated from an inversion-recovery experiment; in all cases, the values were less than 1.5 s, therefore, the inter-transient delay of 8 s (> 5 *T*_1_ values) was used in the EXSY experiments to yield spectra that were suitable for quantitative kinetic analysis.

**FIGURE 2.**
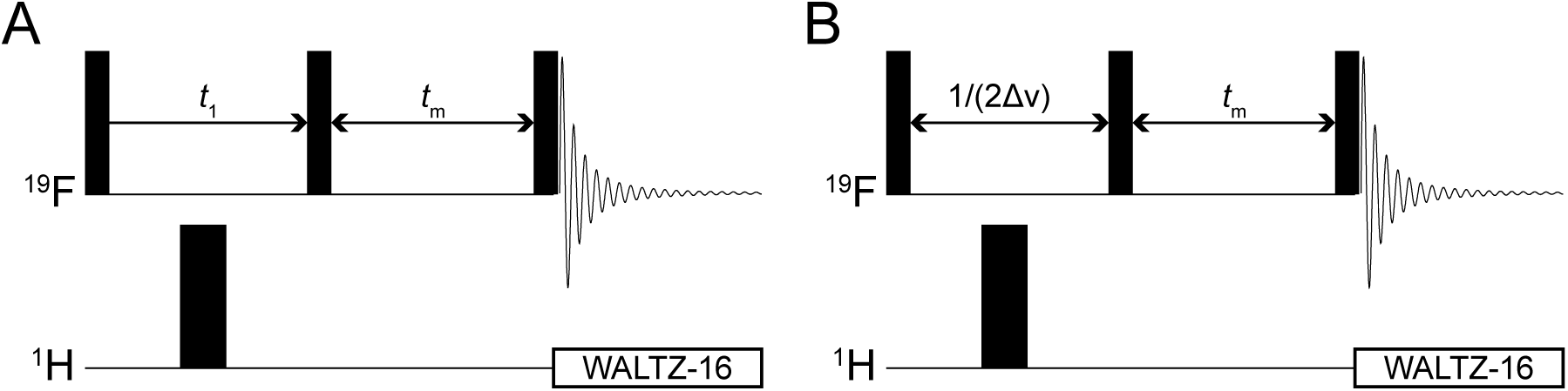
NMR pulse sequences used in the study: (*A*) ^19^F 2D-EXSY with ^1^H decoupling in the two dimensions; (*B*) ^19^F 1D-EXSY: the incremented period (*t*_1_) is replaced by a fixed delay 1/(2Δ*v*), where δ*v* is equal to the difference in the chemical shift between the resonances (in Hz) of the two exchanging populations. Horizontal lines labelled ‘^1^H’ and ‘^19^F’ represent separate radio-frequency (RF) channels tuned to the resonance frequency of ^1^H and ^19^F, respectively. Thin and thick black rectangles represent 90° and 180° hard RF pulses, respectively, applied at the frequency of the corresponding channel. The decaying oscillating curve stands for signal (FID) acquisition, while the open rectangles represent WALTZ-16 composite pulse decoupling (24).

In 2D-EXSY experiments, two transients were accumulated per a free induction decay (FID), in order to suppress axial peaks by a two-step phase cycle of the first 90° pulse (*x*, -*x*; acq: *x*, -*x*) (25). In 1D-EXSY experiments, four or eight transients were acquired per FID (the larger number was used for lower concentrations of FDG), and a four-step phase cycle of the last 90° pulse was used to select the zero-order spin coherence during the mixing time (*x, y*, -*x*, -*y*; acq: *x, y*, -*x*, -*y*). In 2D-EXSY spectra, the mixing time was 0.5 s, while in 1D-EXSY experiments, several mixing times were used (0 – 5 s). The 1D-EXSY spectra were processed with exponential line-broadening of 4 Hz, while the 2D-EXSY spectra were processed with a cosine-squared window multiplication function in both frequency dimensions. The processed NMR spectra were imported into *Mathematica* (Wolfram, Champaign, IL) (26), where peak integration and subsequent numerical data analysis were performed. *Mathematica*’s routine ‘NonlinearModelFit’ was used for nonlinear regression analysis.

## RESULTS

### Stability of RBCs in inosine medium

Accurate membrane transport studies of GLUT1 require that the RBCs are maintained in a stable morphological and metabolic state in a glucose-free medium prior to the experiments (and then in the presence of the glucose analogues during the experiment). Therefore, given the prolonged incubations (up to three hours) used for many of the experiments, we first aimed to establish and verify an optimal glucose-free medium, composed to prolong the integrity of metabolically-active human RBCs during the fluorosugar ^19^F-NMR EXSY experiments.

In order to provide a non-glucose-based energy source that is required for RBC survival over several hours, an inosine-based saline with additional pyruvate, phosphate and KCl was employed. Pyruvate, as the oxidized counterpart of lactate in the lactate dehydrogenase (E.C. 1.1.1.27) reaction, helps to maintain a higher concentration of NADH that promotes flux through the second half of glycolysis, starting with glyceraldehyde-3-phosphate dehydrogenase (E.C. 1.2.1.12) and reduces spontaneous oxidation reactions. KCl provides the extracellular cation for continual activity of Na^+^,K^+^-ATPase, the dominant site of adenosine triphosphate (ATP) hydrolysis in the cell and, therefore, a major determinant of glycolytic flux. Given the known beneficial effects of inosine, pyruvate and KCl, it was decided to incorporate them into a saline with the following final composition: 10 mM inosine; 10 mM sodium pyruvate; 5 mM NaH_2_PO_4_; 4.5 mM NaOH; 10 mM KCl; 133 mM NaCl in Milli-Q H_2_O, which provided osmolality of 300 mOsm kg^-1^ and pH 7.4.

In order to assess the stability of RBCs in the medium, we used ^1^H spin-echo NMR experiments (27) to monitor ongoing production of lactate from glycolysis by RBCs in four different media: (I) ‘normal’ saline (150 mM NaCl + 10 mM KCl); (II) ‘normal’ saline with addition of 10 mM Glc; (III) the ‘standard’ ascorbate saline that was used in some previous ^19^F-NMR studies of fluorosugars with RBCs (21, 28, 29); and (IV) the developed inosine saline. The ^1^H spin-echo NMR spectra of each medium were recorded for 5 min (using spin-echo delay τ = 67 ms) in 1-hour intervals; the obtained spectra are shown as stacked plots in Fig. 3 *A*.

**FIGURE 3.**
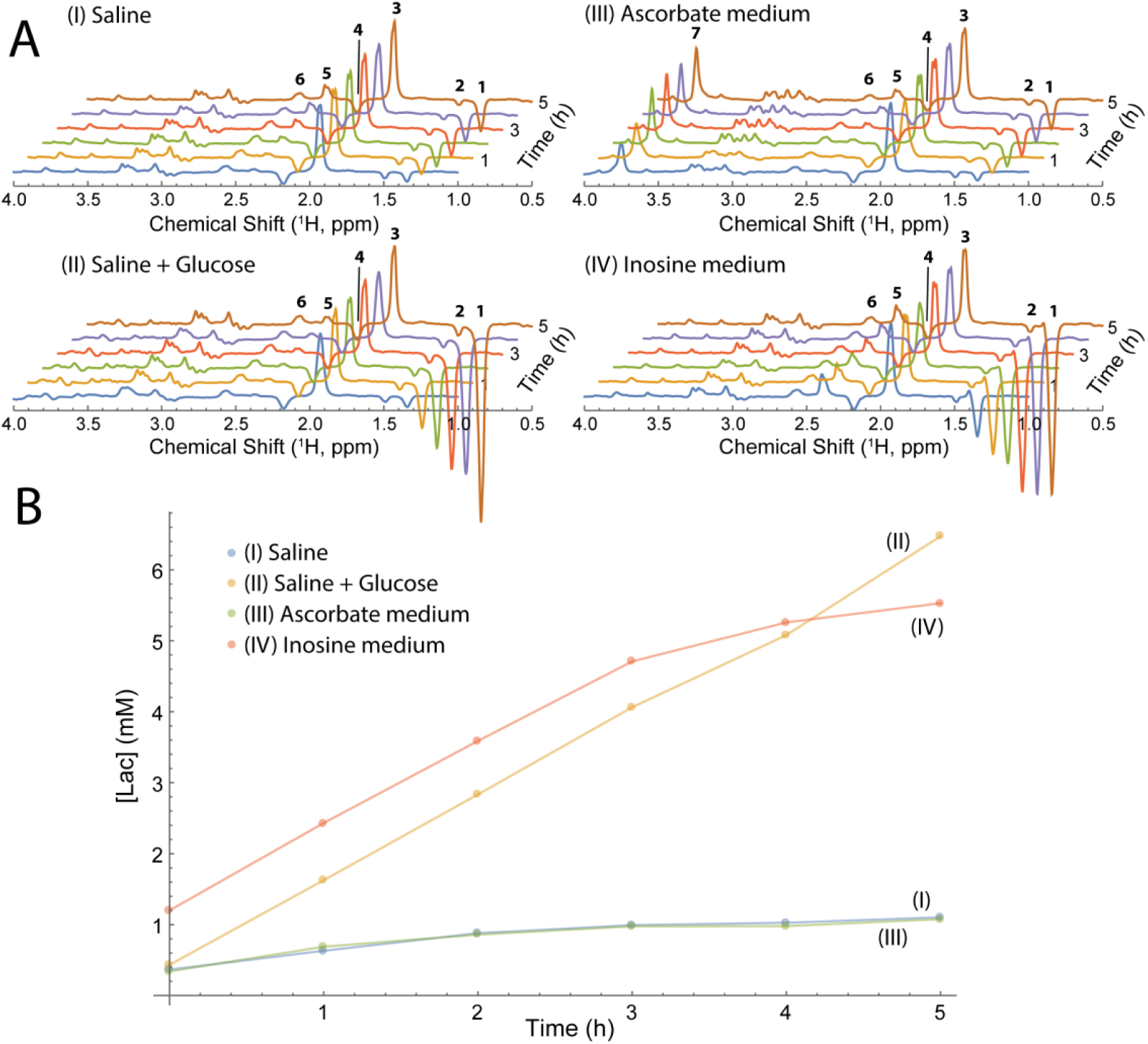
Lactate production by RBCs in four suspension media. (*A*) 400.13 MHz ^1^H NMR spin-echo spectral time courses of RBC suspensions of the four studied media (see text). The following assignments were made by adding individual substances to the samples in a separate experiment and using previously published data (30): **1** (1.34 ppm) – C<u>H</u>_3_ of lactate; **2** (1.49 ppm) – C<u>H</u>_3_ of alanine and/or pyruvate hydrate; **3** (1.93 ppm) – C<u>H</u>_3_ of acetate; **4** (2.18 ppm) – C<u>H</u>_2_(β) of the glutamyl residue in reduced glutathione; **5** (2.39 ppm) – C<u>H</u>_3_ of pyruvate (keto-form); **6** (2.57 ppm) - C<u>H</u>_2_(γ) of the glutamyl residue in reduced glutathione; **7** (3.74 ppm) – Tris. The spectra were scaled so that the intensities of the acetate peak were the same in each series. (*B*) Production of lactate by RBCs, as calculated from the spectra of (*A*), over the duration of the experiment (5 h at 37°C). Coefficients of variation in concentration estimates were uniformly less than 5%.

Six spectral time points were acquired for each of the samples (I-IV) in experiments conducted at 37 °C for 5 h. Each of the samples showed ongoing production of lactate (as a steady growth of peak **1** in Fig. 3 *A*). Figure 3 *B* shows the increase in the lactate concentration over time in the samples in the four different media. The rate of lactate production was significantly greater in the samples containing an energy-supply substrate: glucose (II) and inosine (IV), in comparison with the substrate-free samples: ‘normal’ saline (I) and the ascorbate medium (III) (Fig. 3). We ascribed the production of lactate in samples (I) and (III) to the breakdown of endogenous 2,3-bisphosphoglycerate (23BPG).

In order to quantify the lactate production rates, an internal standard of an inert compound (5 mM acetate) was added to each sample (peak **3** in Fig. 3 *A*). Also, a separate sample with known quantities of acetate (5 mM) and lactate (10 mM) was used to acquire a ^1^H spin-echo NMR spectrum under the same experimental conditions. This allowed us to calculate the coefficients of proportionality, required for the conversion of the measured lactate peak intensities to the corresponding concentrations in mM. Thus, lactate production rates were quantified for the initial linear period of the reaction (0-3 h). There was no statistically significant difference in lactate production rates between sample (II): 1.74 ± 0.03 mmol (L RBC)^-1^ h^-1^ and sample (IV): 1.71 ± 0.03 mmol (L RBC)^-1^ h^-1^, indicating that the glycolytic rate in the inosine-based saline was the same as for the glucose-supplied control. In contrast, samples (I) and (III) did not have any energy-supply substrate and this clearly led to ‘metabolic starvation’ of the cells.

The samples used for the ^1^H spin-echo experiments were also examined with differential interference contrast (DIC) light microscopy. Digital micrographs of each of the samples (in media I-IV) were recorded immediately before the NMR measurements, and after 3 and 5 h of incubation at 37°C; representative micrographs are shown in Fig. 4. The images confirmed the integrity (biconcave-disc shape) of the RBCs in the glucose and inosine media. This indicated that the RBCs survived well and were not adversely affected by any of the solutes and ions in the inosine-saline solution, making it a suitable glucose-free medium for the cells. In contrast, there was a notable formation of RBCs with irregular shapes (early echinocytes) in the samples that did not have energy-supply substrates (these cells are marked with asterisks in Fig. 4).

**FIGURE 4.**
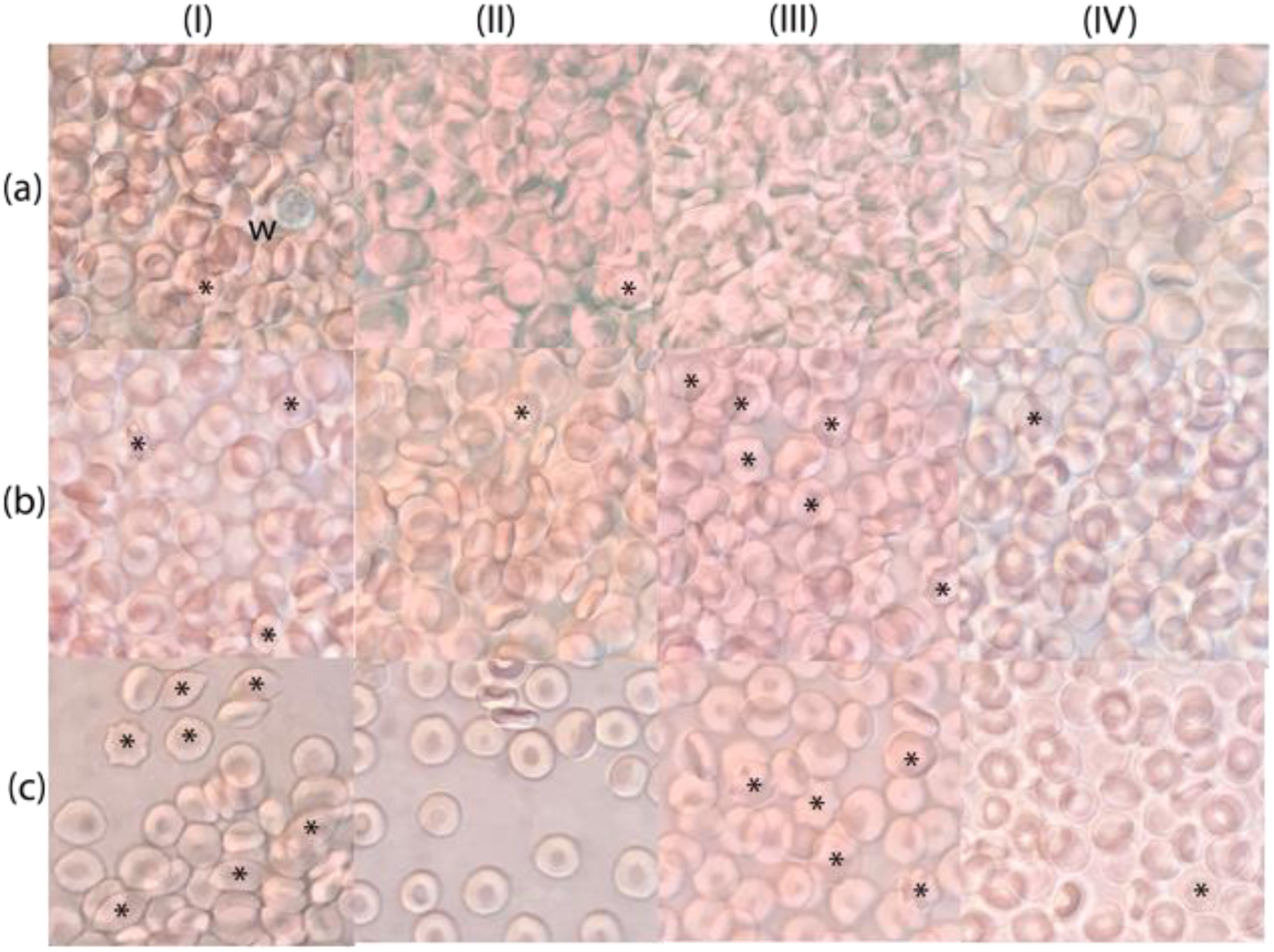
DIC light micrographs of RBCs in samples (in media I-IV) used in the experiment of Fig. 3. The micrographs for each sample were obtained: (a) before the acquisition of the ^1^H spin-echo spectra; and (b) upon 3 h and (c) upon 5 h after the start of the incubation at 37°C. The asterisks indicate the most obvious cells of irregular shape, while ‘w’ labels a white blood cells in micrograph (aI).

Overall, the results indicated that the RBCs were metabolically stable in ‘inosine saline’ for at least 3 h at 37°C. This was sufficient for readily scheduled acquisition of a series of the ^19^F-NMR EXSY experiments, as is described next.

### 2D-EXSY experiments

Initially, we measured 1D ^19^F-NMR spectra of each of the fluorosugars (shown in Fig. added to a suspension of RBCs. The peaks from both anomers present inside and outside the RBCs were clearly visible, and they were assigned using previous reports (20, 21). The mutarotation constant *k*_mut_ = [*β*]/[*α*] and anomeric ratio *a* (Eq. 9) of the three sugars inside and outside the cells were readily measured from the ratio of the peak integrals, and it was verified that *a*_inside_ and *a*_outside_ were equal in all cases (within a ∼3% margin of error). The values of the anomeric ratio *a* for each of the three fluorosugars are given in the first column of Table 2.

2D-EXSY ^19^F-NMR spectra were recorded from the three studied monofluorinated analogues of Glc in RBC suspensions using the pulse sequence of Fig. 2 *A*. These spectra showed no cross-peaks between the spin populations of the α- and β-anomers (Figs. 5-7 for FDG-2, FDG-3 and FDG-4, respectively), as observed by O’Connell et al. (21). Importantly, this indicated that the anomeric exchange (mutarotation) was negligible on the timescale of the EXSY mixing time (0.5 s) and could safely be ignored in the subsequent kinetic analysis. By contrast, for each anomer there were prominent cross-peaks between the resonances of the corresponding extracellular and intracellular spin populations (Figs. 5-7), reflecting the extent of their transmembrane exchange.

**FIGURE 5.**
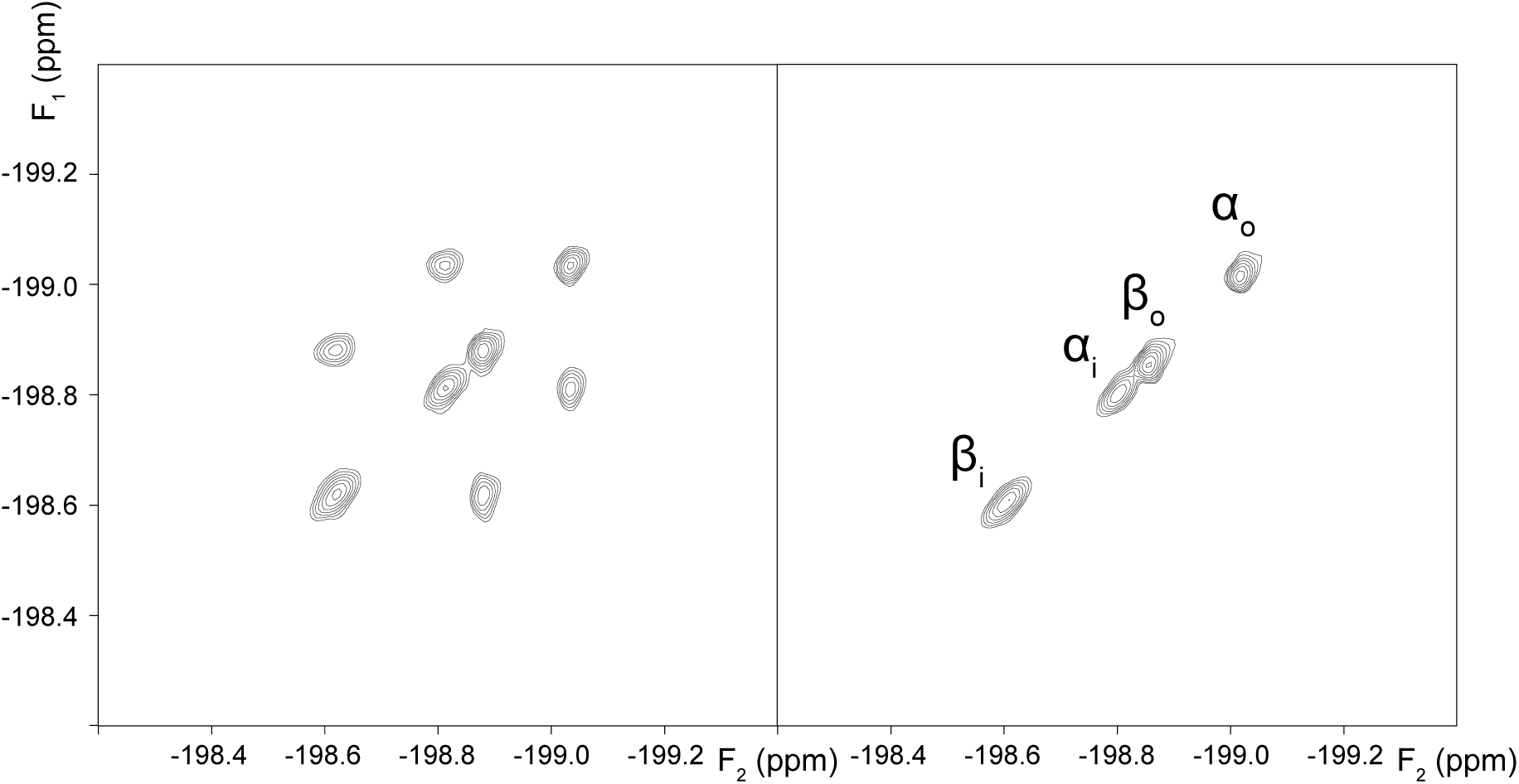
2D-EXSY ^19^F-NMR spectra (376.46 MHz) of 12.4 mM FDG-2 in an RBC suspension in the absence (left panel), and presence of 0.1 mM cytochalasin B (right panel). The spectra were recorded with the pulse sequence shown in Fig. 2 *A*, using a mixing time *t*_*m*_ = 0.5 s.

In a control experiment, it was verified that the exchange cross-peaks completely disappeared in the presence of 0.1 mM cytochalasin B, a known inhibitor of glucose transport in human RBCs (31), as shown in Figs. 5-7. This was consistent with transmembrane exchange of the FDGs being mediated by GLUT1, while the contribution of passive permeation by diffusion was negligible (in the mixing time of the EXSY experiment).

**FIGURE 6.**
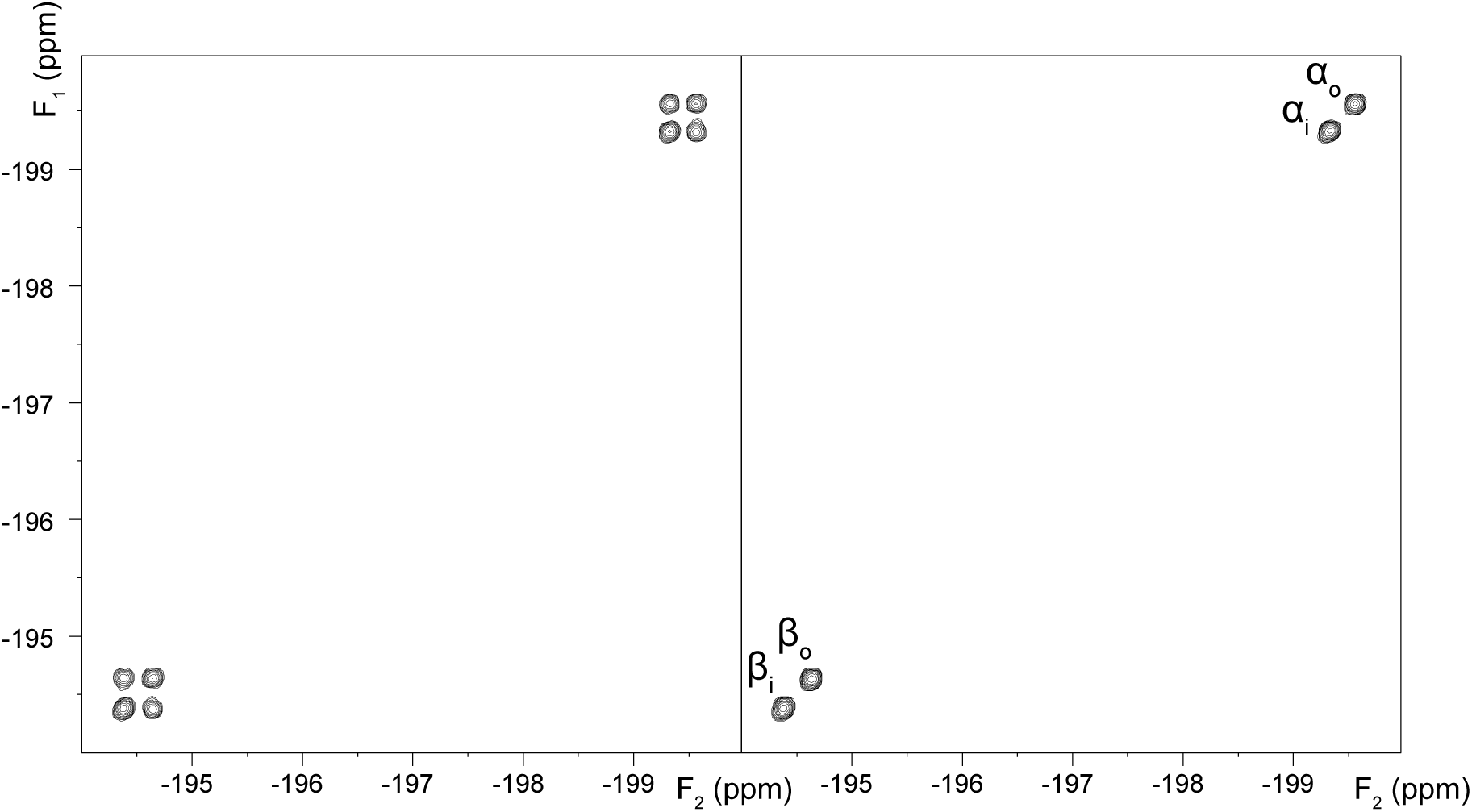
2D-EXSY ^19^F-NMR spectra (376.46 MHz) of 12.4 mM FDG-3 in an RBC suspension in the absence (left panel), and presence of 0.1 mM cytochalasin B (right panel). The spectra were recorded with the pulse sequence shown in Fig. 2 *A*, using a mixing time *t*_m_ = 0.5 s.

**FIGURE 7.**
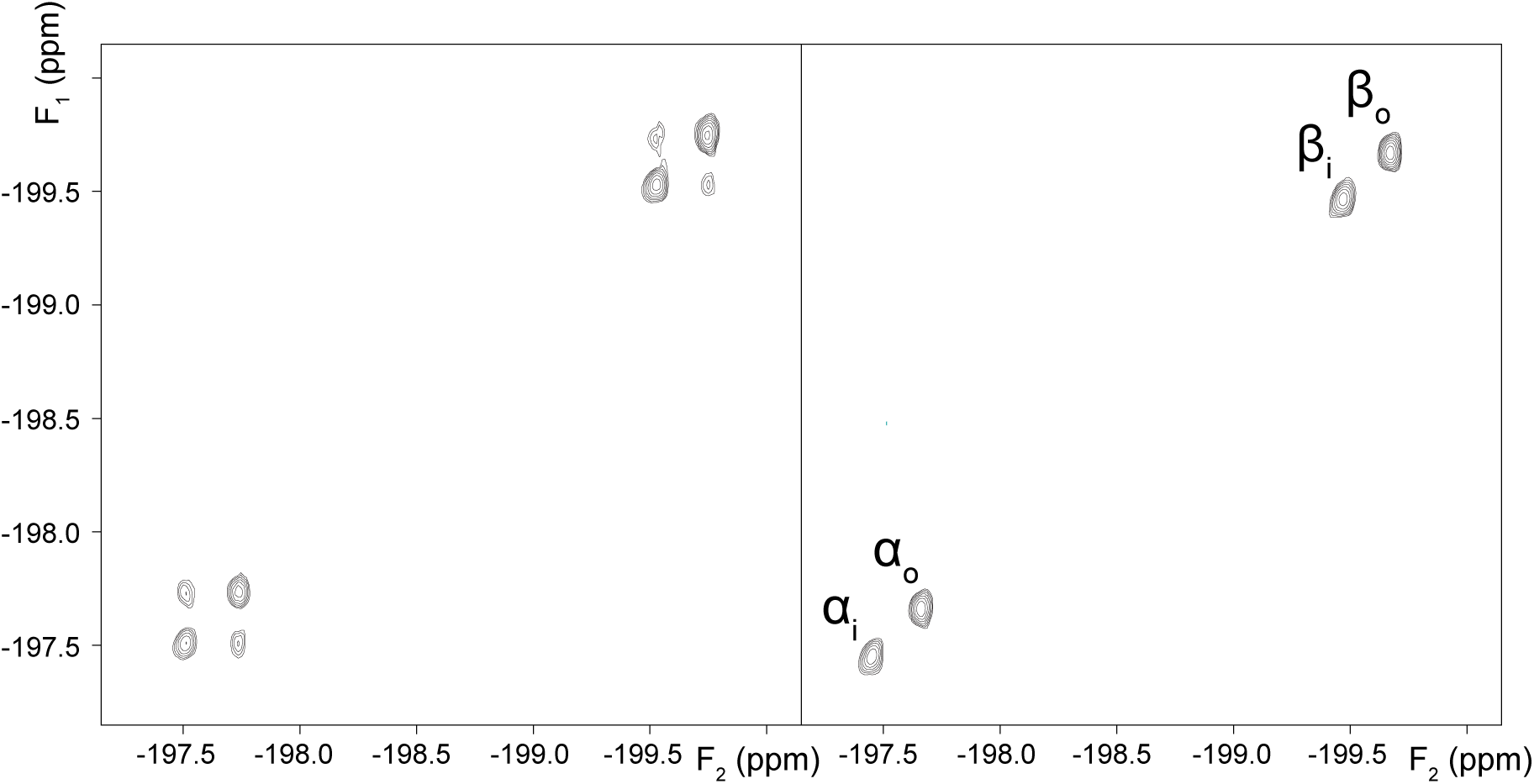
2D-EXSY ^19^F-NMR spectra (376.46 MHz) of 6.2 mM FDG-4 in an RBC suspension in the absence (left panel), and presence of 0.1 mM cytochalasin B (right panel). The spectra were recorded with the pulse sequence shown in Fig. 2 *A*, using a mixing time *t*_m_ = 0.5 s.

### Determination of anomer exchange rates constants

In order to measure the rate constants for influx (*k*_ei_) and efflux (*k*_ie_), the 1D-counterpart of the 2D-EXSY experiment (32) was used. It allowed more precise measurements in a shorter experiment time. From Figs. 5-7, it is clear that, for a given fluorinated sugar, there are two independent pairs of exchanging spin populations. Thus, the rates of magnetization exchange of each anomer can be described by a pair of simultaneous two-site Bloch-McConnell differential equations (33). In the context of the 1D-EXSY experiment, the time dependence of *longitudinal* magnetization of the extracellular *M*_e_(*t*) and intracellular *M*_i_(*t*) spin populations, during the mixing time *t*_*m*_, is given by:

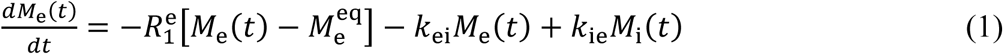

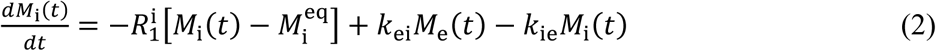

where 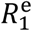 and 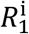 are the longitudinal relaxation-rate constants, characterizing the return of the longitudinal magnetizations to the corresponding thermal-equilibrium values, 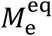 and 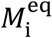. *k*_ei_ and *k*_ie_ are the rate constants for influx and efflux, respectively, as defined by Eqs. S4-S6 in the Supporting Material.

Solution of the Bloch-McConnell differential equations gives the dependence of the intensities of the NMR signals in a magnetization-transfer experiment on the mixing time (34). In the 1D-EXSY experiment, employed for quantification in the present work, the radio-frequency transmitter was set ‘on resonance’ with the intracellular spin-population in order to achieve the selective inversion of its magnetization at the beginning of the experiment (pulse sequence of Fig. 2 *B*); this specified a key initial condition for the solution of the differential equations: 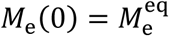 and 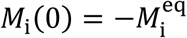 for Eqs. 1-2.

The 1D-EXSY experiment was repeated for 16 different mixing times ranging from 0 to 5 s. The integrals of the two NMR signals were recorded at each mixing time; this generated time-course plots for the two exchanging magnetizations, as functions of the mixing time, *viz*., *M*_e_(*t*_m_) and *M*_i_(*t*_m_). Fitting the numerical solutions of the Bloch-McConnell equations (Eqs. 1-2) to the experimental dependencies of *M*_e_ and *M*_i_ on the mixing time yielded estimates of the rate constants *k*_ei_ and *k*_ie_ for both anomers, together with their standard errors. As described in the Supporting Material, it is more convenient to work with the efflux rate constants *k*_ie_ rather that influx rate constants *k*_ei_ because the former does not dependent on the hematocrit values (21). Due to the saturation of GLUT1 with substrate molecules, the exchange rate constants are functions of substrate concentration. Therefore, we repeated the acquisition and processing of the 1D EXSY spectra at five different substrate concentrations, and the statistically estimated *k*_ie_ values are given in Table 1.

**Table 1.**
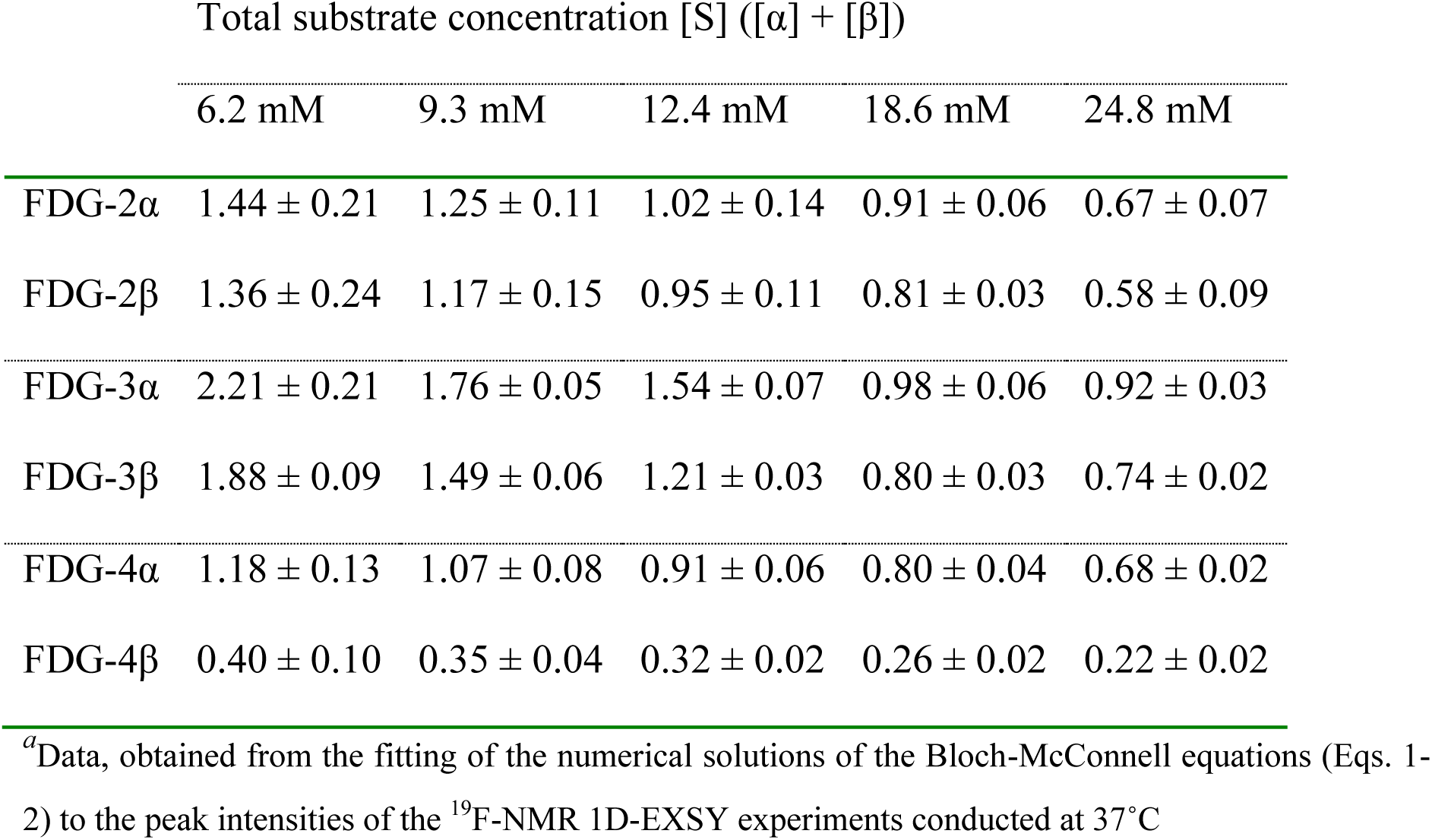
Values of the apparent efflux rate constant *k*_ie_ (s^-1^) at different substrate concentrations^*a*^

Next, we used the values of *k*_ie_ for the calculation of the equilibrium-exchange rate *v* of each anomer (by combining Eq. S4 with Eq. S14):

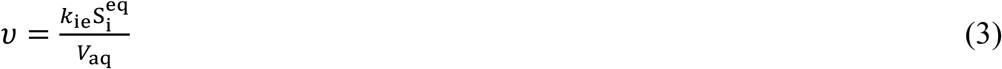

where 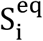 is the mole quantity of the substrate in the intracellular compartment at equilibrium; and *V*_aq_ is the total aqeuous volume of the sample. It is convenient to normalize the transport rate *v* with respect to the total concentration of the carrier and substrate in the sample. This leads to the following expression:

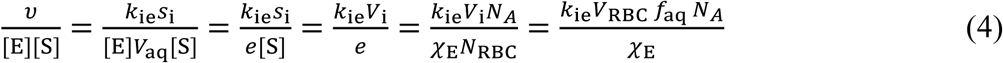

where ***e*** is the total mole quantity of the carrier in the sample; *χ*_E_ is the number of copies of the enzyme per cell (∼200,000 for GLUT1 (35)); ***N***_RBC_ is the total number of cells in the sample; *V*_i_ is the total intracellular volume of the sample, accessible to solutes; *V*_RBC_ is the average physical volume of one RBC (86 fL in isotonic solution (23)); *f*_aq_ = 0.717 is the fraction of the intracellular volume that is aqueous (36); and ***N***_A_ = 6.02214 × 10^23^ mol^-1^ is Avogadro’s number. From Eq. 4, it is evident that the normalized transport rate is directly proportional to the efflux rate constant *k*_ie_ and is independent of *Ht*. Nevertheless, because of the saturability of the carrier by substrate, the exchange rate constants *k*_ei_ and *k*_ie_ (and transport rates *v*) are nonlinear functions of the respective anomer concentrations (Table 1).

### Determination of the catalytic activity of GLUT1 on individual anomers

The equilibrium-exchange kinetics of GLUT1-mediated transport can be described by Michaelis-Menten steady-state enzyme kinetic equations that have been extended to account for competing substrates (see Supporting Material). The overall reaction scheme for transmembrane-exchange of the two anomeric species is shown in Fig. S3. We developed the equations for equilibrium-exchange kinetics in the most general form, avoiding previously used assumptions about the equality of any of the Michaelis-Menten parameters for the two anomers (20, 37). Namely, we showed that the simultaneous presence of α- and β-anomers of a sugar that are competing for transport by a carrier is accounted for by introducing factors 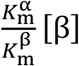 and 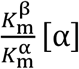 in the denominator of the classical Michaelis-Menten equations (Eqs. S108-S109):

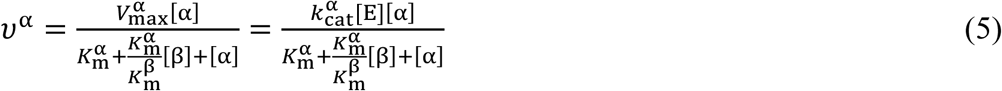

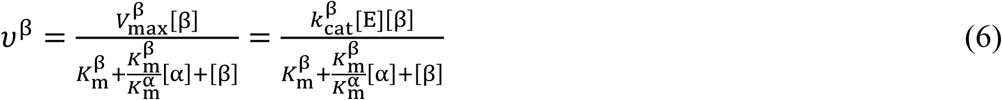

where [E] is the total molar concentration of the enzyme/carrier. *V*_max_, *k*_m_ and *k*_cat_ have the usual meanings, *viz*., the maximum velocity, Michaelis constant and the turnover number, respectively. Superscripts α and β denote the individual values of *V*_max_, *k*_m_, and *k*_cat_ that are specific for each anomer under equilibrium-exchange conditions. *v*^α^ and *v*^β^ are the equilibrium-exchange rates of membrane transport. It is more convenient to express Eqs. 5-6 in terms of normalized rates of transport, which could be readily calculated from *k*_ie_ values using Eq. 4:

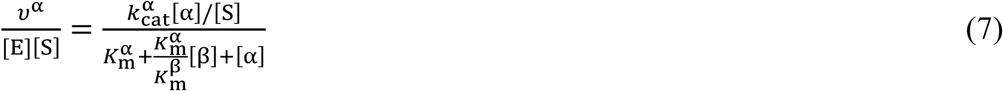

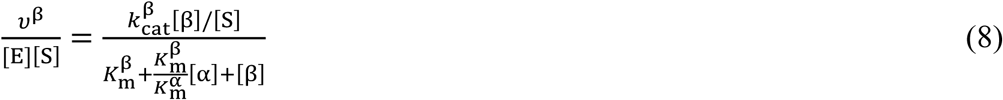

As established earlier using 1D ^19^F-NMR, at equilibrium, the ratio of the concentrations of the two anomers in each compartment is constant and can be defined by the parameter *a*:

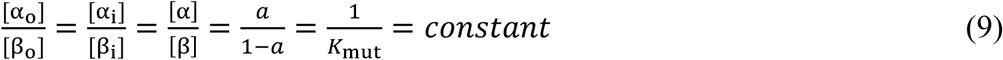

Making the substitutions [α] = *a*[S] and [β] = (1 − *a*)[S] into Eqs. 7-8 gives (also see the equivalent Eqs. S111-S112 in the Supporting Material):

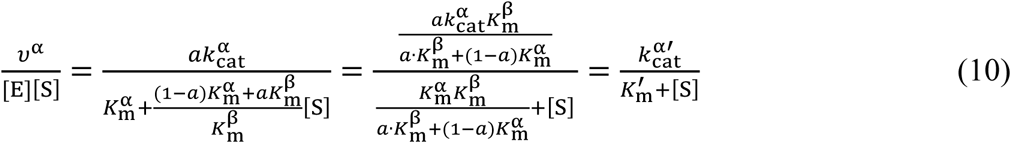

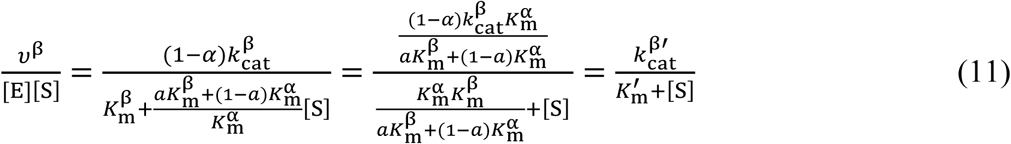

In Eqs. 10-11, the following definitions are made in order to obtain the ‘apparent’ Michaelis-Menten equations:

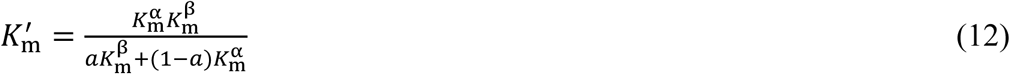

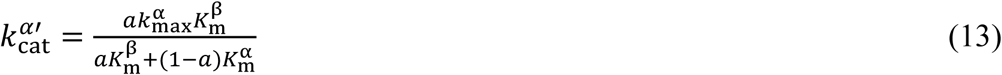

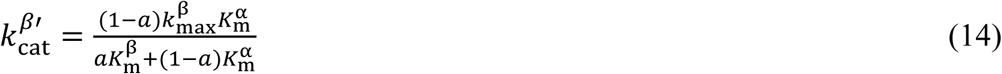

The *specificity constant k*_cat_/*K*_m_ is an important measure of the catalytic efficiency of an enzyme on a particular substrate (1). In the context of carrier-mediated facilitated diffusion, *k*_cat_/*K*_m_ is equivalent to the rate constant of the transmembrane exchange when [S] ≪ *K*_m_ and can be interpreted as the overall membrane permeability of the substrate (38). Unlike the apparent rate constants and transport rates, the overall membrane permeability, defined as *V*_max_/*K*_m_ (or normalized by the amount of enzyme in the medium and expressed as *k*_cat_/*K*_m_), does not depend on the substrate concentration. Additionally, the value of *V*_max_/*K*_m_ is the same for the equilibrium-exchange, zero-trans influx and zero-trans efflux kinetics, which makes it an important characteristic of a carrier-mediated membrane transporter (38).

From Eqs. 12-14, it follows that we can calculate the individual specificity constants as (see Eqs. S118-S119 for derivation):

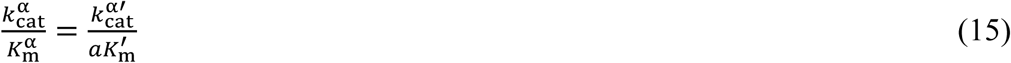

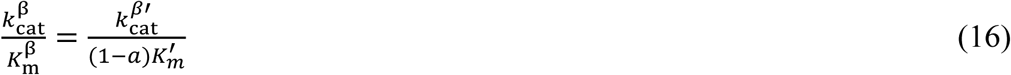

Thus, we proceeded to fit the model of Eqs. 10-11 to the obtained dependencies of *k*_ie_ (converted to the values of 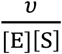 using Eq. 4) on the total substrate concentration [S] (Table 1). This procedure yielded values of 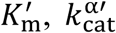 and 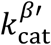 (listed in Table 2), which allowed estimation of the individual specificity constants of GLUT1 towards the two anomers (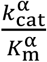 and 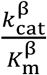) by using Eqs 15-16. The determined *k*_cat_/*k*_m_ values are listed in the last column of Table 2.

**Table 2.** Values of the Michaelis-Menten parameters of GLUT1 for the studied fluorinated sugars, at 37°C

| | $a^a$ | $K_m', \text{mM}^b$ | $k_{\text{cat}}', \text{s}^{-1}b$ | $k_{\text{cat}}/K_m, 10^5 \times \text{M}^{-1} \text{s}^{-1}c$ |
| --- | --- | --- | --- | --- |
| FDG-2α | $0.45 \pm 0.02$ | $10.5 \pm 1.6$ | $4490 \pm 350$ | $9.5 \pm 1.7$ |
| FDG-2β | | | $4150 \pm 330$ | $7.2 \pm 1.3$ |
| FDG-3α | $0.46 \pm 0.01$ | $4.6 \pm 0.9$ | $4490 \pm 290$ | $21.2 \pm 4.4$ |
| FDG-3β | | | $3730 \pm 250$ | $15.0 \pm 3.1$ |
| FDG-4α | $0.42 \pm 0.02$ | $18.5 \pm 1.3$ | $5430 \pm 240$ | $6.9 \pm 0.6$ |
| FDG-4β | | | $1820 \pm 90$ | $1.7 \pm 0.2$ |

<sup>a</sup>Values, determined experimentally from the ratio of the integrals of the resonances in 1D $^{19}\text{F}$ -NMR spectra (Eq. 9)
<sup>b</sup>Values, obtained from fitting of the solutions of the Eqs. 10-11 to the determined efflux rate constants (Table 1)
<sup>c</sup>Values, calculated from the determined values of $a$ , $K'_m$ , $k_{cat}^{\alpha'}$ and $k_{cat}^{\beta'}$ using Eqs. 15-16

## DISCUSSION

### Stability of RBCs in inosine medium

RBCs require a constant supply of energy in their suspension medium to maintain their shape and regulated oxygen affinity. The energy is largely dissipated in the regeneration of ATP, the cell’s ‘energy currency’, from inorganic phosphate and adenosine diphosphate. *In vivo*, the natural energy-supply substrate for human RBCs is Glc (in the blood plasma). As a consequence, during *in vitro* experiments on RBCs, it is usual to add glucose as a metabolic substrate for maintenance of physiological ATP levels. However, since Glc would inevitably interact with GLUT1 during the fluorosugar NMR experiments, it was important to remove all endogenous glucose in the RBC preparations. Under such conditions, only 23BPG remains as an energy source for the RBCs (originally at ∼5 mM), and this normally lasts for only 1-2 h at 37°C, at which point the cells begin to convert to echinocytes (39, 40). Hence, an alternative energy substrate was required for the prolonged EXSY experiments.

In a previous RBC study of FDG-3, Potts and Kuchel used 10 mM inosine in phosphate-buffered saline (pH 7.4) (20). This purine nucleoside is transported by a specific nucleoside transporter in RBCs (41). Inosine undergoes phosphorolysis via purine nucleoside phosphorylase (E.C. 2.4.2.1) to yield ribose 1-phosphate, obviating the requirement for ATP hydrolysis to phosphorylate the saccharide. Then, the constituent atoms enter glycolysis via the pentose phosphate pathway (42). The use of 5 mM ascorbic acid (vitamin C; ascorbate at physiological pH) in Tris-HEPES-buffered saline has also been used as a medium for fluorosugar NMR experiments (21, 28, 29). However, this is not optimal, because ascorbate, if oxidized by dissolved O_2_, is converted to dehydroascorbate (DHA), which is known to be transported via GLUT1 (43, 44). Thus, it might potentially interfere with fluoroglucose transport. In fact, it was recently shown that the binding of Glc by GLUT3 is substantially reduced in the presence of DHA (14).

Moreover, as is clear from the results in Fig. 3, ascorbic acid is not metabolized in human RBCs, so it does not act as a source of free energy. Nevertheless, it may play a role in the reduction of α-tocopherol in the cell membrane (45). We surmise that the primary role of ascorbate in the Glc-free media (21, 28, 29) is to spontaneously transfer electrons to oxidized glutathione and NAD. In turn, NADH transfers electrons to Fe(III) in methemoglobin thus keeping most of the hemoglobin in the RBCs in a diamagnetic state; this is optimal for the signal-to-noise and resolution in NMR spectra. Glc acts as both a source of free energy for the regeneration of ATP, and as a reducing agent via its contribution to the conversion of NAD to NADH and NADP to NADPH. Thus, ascorbate is not required in a medium which contains an energy source that feeds into glycolysis at the glyceraldehyde-3-phosphate dehydrogenase step.

Tris is known to react with aldehydes of low molecular weight in aqueous solution, so it interacts with glyceraldehyde 3-phosphate (46), and potentially with the open-chain form of glucose and its fluorinated analogues. With this in mind, we avoided the ascorbate/Tris-based medium, and developed a different buffer for the ^19^F-NMR EXSY experiments; this was done by combining the known metabolically beneficial properties of inosine, pyruvate and KCl. The results of Figs. 3-4 proved the stability of the RBCs under the new conditions for a prolonged time (at least 3 h at 37 °C).

### Transmembrane-exchange kinetics

Membrane transport of Glc in RBCs is sufficiently rapid for equilibrium across the compartments to be achieved within a few seconds, and yet the transmembrane-exchange at equilibrium is slow on the NMR timescale. In 1D ^19^F-NMR spectra of FDGs in RBC suspensions, this is highlighted by the well-resolved resonances from the spin populations inside and outside the cells (19, 21). This fortuitous property, referred to as a ‘split peak effect’, enables the use of magnetization-exchange spectroscopy (EXSY), for the quantification of the kinetics of the transmembrane exchange (20-22, 47). Furthermore, the α- and β-anomers of these sugars have different ^19^F-NMR chemical shifts thus allowing an examination of the differences in the transport rates of the two anomers (19-21).

In the past, transport of fluorinated sugars has been described by permeability coefficients *P* (cm s^-1^), or apparent exchange rate constants *k* (s^-1^), which can be measured for each anomer. However, these values are functions of the solute concentration, since the kinetics of the transmembrane exchange of Glc and its analogues are well described by Michaelis-Menten enzyme kinetics, with a steady state of the concentration of the carrier-Glc complex (48). Thus, the transport kinetics of a given monosaccharide should ideally be described by estimating the individual Michaelis-Menten parameters (*k*_m_, *V*_max_, *k*_cat_) for each of the two anomers. Since both anomers of a monosaccharide simultaneously compete for binding in the carrier active site (49, 50), there are additional theoretical complications in the kinetic theory (see Supporting Material). Explicit modifications of the rate equations are required and a general form of these has been worked out (37, 48).

Previous anomer-specific studies of glucose analogues in RBCs have relied on assumptions, such as equality of the Michaelis constants for binding to GLUT1 for the α- and β-anomers. This enabled the authors to simplify their rate equations and attribute any differences in the anomeric transport rates to differences in the apparent *V*_max_ (*or k*_cat_) values (20). However, the assumption of equal affinity is not generally valid, since the recent crystal-structure data show that the anomeric -OH group can directly participate in hydrogen bonding inside the active site of the transporter (14). Moreover, a single change in the configuration of one of the carbon atoms might lead to a significant change in the affinity of a monosaccharide for GLUT1. *E.g*., the Michaelis constant of galactose (C-4 epimer of glucose) for GLUT1 is ∼10 times greater than that of glucose (51, 52). This highlights the importance of anomer-specific kinetic studies and implies that *k*_m_and *V*_max_for monosaccharide transport, reported in the literature as a single or common value, might not reflect the true complexity of the transport kinetics of the GLUTs (53).

Previously reported anomer-specific data on the transport of glucose via GLUT1 were contradictory. An early study by Faust concluded that the β-anomer of Glc penetrates the RBC membrane ∼3 times faster than the α-anomer (54). Miwa et al. reported ∼1.5 times faster influx of the β-anomer into cells, but the same rate for the efflux of the two anomers (55). This report was consistent with the view of Barnett et al. (56) that the C-1 hydroxyl group of Glc interacts with GLUT1 only on influx, but not on efflux.

Carruthers and Melchior reported no difference in transport rates and affinity by the glucose transporter of the two anomers of Glc, by using inhibition-uptake studies of radioactive Glc (49) in intact RBCs. This result was confirmed in a more recent study by the same group, and they concluded that GLUT1 does not prefer any specific anomer of Glc (37) in RBC ghosts.

Other reports have indicated that the β-anomer is favored at higher temperatures, while the α-anomer is preferred at lower temperatures (50) and that α-anomer is 37% more efficient in promoting the GLUT1 conformational change than the β-anomer (57). Using a similar methodology to the present study (equilibrium-exchange magnetization-transfer experiments), Kuchel et al. measured the transport rates of Glc at 40°C and concluded that the transport of the α-anomer was ∼1.7 times faster than that of the β-anomer (58).

London and Gabel used fluorine-substituted at position C-1 analogues of glucose in order to halt the mutarotation between the two anomeric species (47). Thus, two 1-fluoro-1-deoxy-D-glucose (FDG-1) analogues of glucose were used in two separate experiments in order to probe their transport rates. Only the β-anomer of FDG-1 was transported under equilibrium-exchange conditions (47); while an earlier study by Barnett et al. reported that the entry of sorbose into RBCs was inhibited by both anomers of FDG-1, with inhibition constants of ∼15 mM (β) and ∼80 mM (α) (59).

These inconsistent results in the literature are probably due to the complicated mechanism of transport by GLUT1 (53): it involves several conformational changes of the protein as reported in the recent structural studies (13, 14). Thus, the individual anomers might be more efficient at each of the given transport/binding steps, which complicates the comparison between the anomer-specific kinetic rates. Specifically, the overall transport rates would depend on the conditions of the experiment and the choice of the measurement-assay parameters. Furthermore, the kinetic rate constants and the values of *V*_max_/*k*_cat_/*k*_m_ would be different for equilibrium-exchange, zero-trans, zero-cis, infinite-trans and infinite-cis experiments (38, 48, 51). Additional complications arise from the fact that the previously reported experiments were different with respect to the measurement mode: some of them employed direct quantification of the transport rates of monosaccharides, while others used an inhibition type of experiment, by measuring the influx/efflux of other competing metabolites. More importantly, however, the transport rates in their own right are not sufficient for the discrimination of the GLUT1 efficiency in transporting various Glc analogues and their anomers, since, as discussed above, these values are dependent on the actual substrate concentration used in the experiments.

With this in view, we propose the use of the simple NMR-based assay, as described in the current work, to discriminate the catalytic efficiency of GLUT1 for transport of various fluorine-labelled monosaccharides by measuring the *k*_cat_/*k*_m_ values, obtained in equilibrium-exchange ^19^F-NMR EXSY experiments. The advantage of the proposed approach is that these values can be obtained simultaneously for both anomers of a monosaccharide without requiring separation of the two anomeric species (which is not possible in aqueous media) or solution compartments. In addition, *k*_cat_/*k*_m_ values are equal for the equilibrium-exchange, zero-trans efflux and zero-trans influx experiments, thus accounting for a range of conditions under which they could be measured and compared for different sugar anomers (38, 51). Thus, our approach allows more facile comparison of the GLUT1 catalytic efficiency on different substrates, including anomeric species of the same sugar.

Overall, the efflux rate constants obtained in the present study (Table 1) agree well with previously reported values for the same or related sugars (20, 21). Our results given in Table 1 confirm that the apparent efflux rate constants decrease with increasing substrate concentration, that the α-anomer is transported faster than the β-anomer for these sugars, and that FDG-3 is transported faster than FDG-2, which in turn is transported faster than FDG-4 (21).

For FDG-3, we estimated slightly higher rate constants than reported by O’Connell et al. (21). We postulate that this was due to the presence of 5 mM ascorbate in the medium used previously by these researchers. Indeed, our rate constants for FDG-3 are more similar to those reported by Potts and Kuchel, using experiments which were carried out in the presence of inosine (but no pyruvate), rather than ascorbate (20). Nevertheless, Dickinson et al. (22) reported even higher efflux rate constants for FDG-3; their values were in the range 2.12 – 2.40 s^-1^ (β) and 1.99 – 2.48 s^-1^ (α) (obtained at a substrate concentration of 10 mM). As the authors indicated, such high values could be due to the fact that all the blood donors were pregnant women and the RBCs might have been atypical.

It was also clear that the variations in transport rate with increasing sugar concentration were not equivalent for each sugar (*e.g*., FDG-4α was transported more slowly than FDG-2α at a concentration of 6.2 mM, but at an equal rate at a concentration of 24.8 mM). This clearly complicates a structural interpretation of these data; however, an easier comparison between the permeability of various sugars/anomers is done via the estimated specificity constants.

From the data of Table 2, it was evident that variation in specificity constants (*k*_cat_/*k*_m_) of GLUT1 for the different fluorosugars, for the most part, followed the patterns in the concentration-dependence data in Table 1. Specifically, the specificity constant was substantially larger for FDG-3 than for FDG-2, which in turn was larger than for FDG-4 (*t*-test *P* < 0.01). Interestingly, while the *k*_cat_/*k*_m_ value of the α-anomer of FDG-3 was twice that of FDG-2α, and three times that of FDG-4α, the catalytic efficiency of GLUT1 for the β-anomer of FDG-3 was also twice that of FDG-2β, but almost nine times that of FDG-4β. In addition, for each fluorosugar, GLUT1 had a higher specificity constant for the α-anomer than the β-anomer. As noted below, this is surmised to be a consequence of the anomeric preference of binding by GLUT1, which was previously interpreted in terms of a simple alternating conformation model (21, 47).

### Interpretation of kinetic data in the context of GLUT1 structure

We can consider the specificity constants *k*_cat_/*k*_m_ as indicators of the catalytic efficiency of GLUT1 towards a particular anomeric species. As discussed by Percival and Withers, while *k*_m_ values are not reliable indicators of ground-state affinity, the values of *V*_max_/*k*_m_ (or, equivalently, *k*_cat_/*k*_m_) are more easily interpreted, as they reflect the activation-energy barrier of the transition state, with changes indicating the variation in binding interactions between different substrates (60). In the context of membrane transport, the specificity constants are representative of binding at the rate-determining step in the overall transport mechanism (in terms of elementary steps, this incorporates the binding event, the transport event, and the release event, with the transport event consisting of multiple steps in GLUT1 conformational reorganization (14)).

The recent high-resolution crystal structure of GLUT3 protein with bound Glc (PDB code 4ZW9 (14)) can be used to interpret the chemical forces governing the binding of substrates to GLUT-like proteins. GLUT3 is the main transporter of Glc in neurons and shares 64.4% identity with GLUT1 (61). As all the amino-acid residues that are responsible for the binding of Glc are invariant between GLUT1 and GLUT3, we used the residue numbering of GLUT3 below, to be consistent with the original paper (14). In this analysis, GLUT3 was selected over GLUT1 because its structure is of higher resolution. Also, GLUT3 had pure Glc bound, while GLUT1 was bound with *n*-nonyl-β-D-glucopyranoside.

According to Deng et al., the hydrogen bonds formed in the binding of both glucose anomers involve the same residues, including those to the anomeric -OH group (14). The authors indicate that the α-anomer was more abundant (∼69%) in the crystallized protein than the β-anomer, despite the fact that α-form is the minor species of Glc in solution (∼36%). While the corresponding anomeric ratio of binding of the fluorosugars to GLUT1 is not known, the larger *k*_cat_/*k*_m_ values of the three FDG α-anomers, obtained in the present work, also indicate their stronger stabilization of binding in the transport transition state, compared to the respective β-anomers.

The structural data provided by Deng et al. allow deliberation on the impact of deoxyfluorination at the 2-, 3-, and 4-positions of the glucose scaffold. Both Glc anomers were shown to form hydrogen bonds between C2-OH, C3-OH and C4-OH groups and the protein, involving the same amino-acid residues in the protein’s active site for each anomer (14). Therefore, in the following, we consider the binding of the F-substituted moieties without considering the differences between the two anomers.

Specifically, as described by Deng, C4-OH serves as a hydrogen-bond donor with the carbonyl group of Asn286; C3-OH serves as a hydrogen-bond acceptor with the amide side-chains of Asn286 and Gln281; and finally, C2-OH is both a hydrogen-bond donor with the carbonyl of Gln280 and a hydrogen-bond acceptor with the amine of Trp386. These interactions are shown schematically in Figure 8.

**FIGURE 8.**
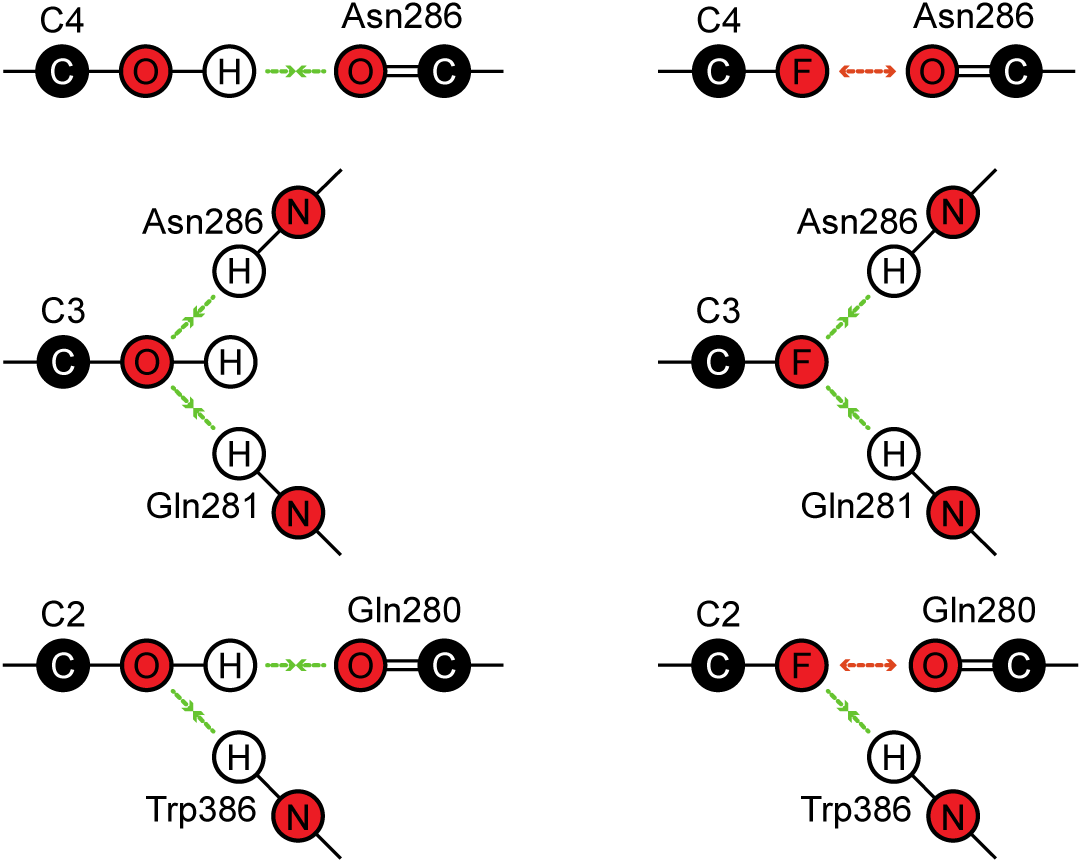
Participation of the -OH groups of Glc in binding to GLUT3 (and by analogy, with GLUT1), according to the PDB structure 4ZW9 (14) (see text).

If the –OH of the C4-OH moiety is substituted by a fluorine atom, an equivalent hydrogen bond cannot be formed, and electrostatic repulsion may exist between the oxygen atom of Asn286 and the F atom of FDG-4 (although the F atom in FDG-4 could also have an attractive interaction with the carbonyl dipole of Asn286 (62)). In contrast, when the –OH of C3-OH is substituted by F, it can still form two hydrogen bonds, potentially leading to a higher apparent binding affinity of FDG-3 in comparison with FDG-4. In the case of FDG-2, there is an intermediate situation upon substitution of –OH by -F, since the C2-OH serves as both a hydrogen-bond donor and acceptor. Therefore, the binding affinity of FDG-2 will be lower than FDG-3, but higher than FDG-4. Although these considerations entail ground-state binding, the obtained *k*_cat_/*k*_m_ values of the three monodeoxyfluorinated glucoses (Tables 1-2) are nevertheless consistent with this picture.

## CONCLUSIONS

We extended the steady-state Michaelis-Menten theory to be applicable for the case when two competing molecular species bind to and cross the cell membrane via the same carrier protein. This novel theory was used to determine the values of the kinetic parameters for three different monofluorinated glucoses (FDG-2, FDG-3 and FDG-4, Fig. 1), as substrates of GLUT1.

To our knowledge, we have shown for the first time that the individual specificity constants of a transporter protein for two anomeric species can be determined from the equilibrium-exchange experiments, without physical separation of the anomers. The experimental approach entails measuring the apparent efflux rate constants of the monosaccharide substrates at several different concentrations. The theoretical development involved extension of the Michaelis-Menten equations for description of the membrane-transport kinetics when two anomeric species of the same compound are in competition for interaction with the carrier.

The developed method was used to quantify RBC membrane transport of the anomers of three mono-fluorinated analogues of glucose in a modified medium, designed to provide stable metabolic properties of the cells over the lifetimes of the experiments. This allowed us to evaluate the specific interaction (specificity constants) of the carrier individually for both anomers of each of the studied fluorosugars. The estimated specificity constants showed faster transport of the α-anomer versus the β-anomer for all investigated substrates, with FDG-4 featuring the largest anomeric preference. A qualitative interpretation of the differences in the transmembrane-exchange rates of FDG-2, FDG-3 and FDG-4 in terms of the perturbation of binding of these monosaccharides to GLUT1, caused by specific F/OH substitutions, has been provided.

Our developed approach and mathematical analysis could become important for a variety of applications such as drug development and systems biology. In turn, such studies can be used to probe responses of membrane proteins in their native environment, *e.g*., in testing whether disease conditions lead to altered permeability of particular carbohydrates, or to changes in cell-membrane integrity (22). Fluorosugars are also of great interest as imaging agents. Radiolabeled 2-fluoro-2-deoxy-D-glucose (FDG-2) is currently the most used PET-imaging substrate for cancer diagnosis (63), while fluorinated fructose analogues have recently been shown to hold promise as imaging agents for GLUT5 expression in breast cancer tissue (64).

## AUTHOR CONTRIBUTION

D.S., B.L. and P.W.K. designed the study. D.S., C.Q.F. and P.W.K. performed the experiments. D.S., I.K. and P.W.K. analyzed the data. D.S. and P.W.K. developed the theory. D.S., B.L. and P.W.K. prepared the manuscript.

## ACKNOWLEDGEMENTS

We thank Dr Ann Kwan for the maintenance of the NMR spectrometer and valuable discussions on setting it up our experiments.

## SUPPORTING CITATIONS

References (65-74) appear in the Supporting Material.

## Supporting Material

### Introduction

We have recently reviewed the techniques relevant to the analysis of equilibrium-exchange magnetization-transfer experiments in studies of membrane transport in cellular systems (1). Special emphasis was put on the differences between notions of transmembrane fluxes, rate constants and permeability coefficients, in the context of these experiments. These differences are briefly revisited here, along with some other important concepts pertinent to the quantification of the transmembrane-transport kinetics in the special case of a carrier-mediated (saturable) transport of two competing species in cell suspensions, with special considerations given to experiments on red blood cells (RBCs).

### Concepts in transmembrane solute-flux analysis

#### Aqueous volume of the sample

When analyzing transmembrane-exchange kinetic data in cell samples, it is important to take into account the fact that the total aqueous volume of the sample, *V*_aq_, is not the same as the total volume of the sample, *V*_tot_ (1). In fact, these two parameters are related to each other in the following way:

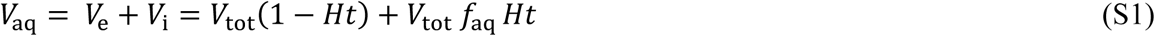

where *Ht* is the value of hematocrit (fraction of the total volume of the sample occupied by RBCs), which is usually known from the sample preparation; *V*_e_ is the total extracellular aqueous volume of the sample; *V*_i_ is the internal aqueous volume that is determined by the coefficient *f*_aq_ = 0.717, which is the fraction of the RBC volume that is taken up by the intracellular water (2).

#### Transmembrane fluxes and rate constants

Transmembrane flux is defined as the mole quantity of the substrate that crosses the membrane per unit of time:

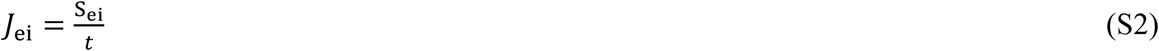

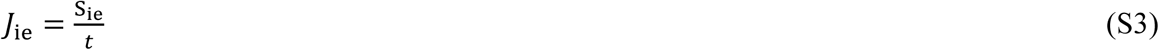

where *J*_ei_ is the substrate flux from outside into the cells (influx; ‘ei’ denotes extracellular-to-intracellular), while *J*_ie_ is the flux in the opposite direction (efflux; ‘ie’ denotes intracellular-to-extracellular). S_ei_ and S_ie_ are the mole quantities of the substrate that are transported in the two directions during the time *t*.

At transmembrane equilibrium, the fluxes in both directions are the same 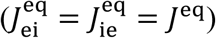 and the apparent rate constants for transport are defined as (1):

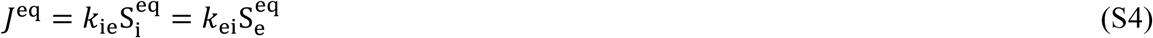

where *k*_ie_ is the apparent rate constant for efflux, *k*_ei_ is the apparent rate constant for influx; *J*^eq^ is the flux of substrate in both directions at equilibrium (units; mol s^-1^); 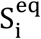 and 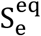 are the equilibrium mole quantities of the substrate inside and outside the cells, respectively. Note that, in order to obtain the flux values, *k*_ie_ and *k*_ei_ are multiplied by the mole *amounts* of the substrates (and not their concentrations); this is a notable difference when a medium contains whole cells (erythrocytes), which introduces compartmentalization. *k*_ie_ and *k*_ei_ are measured in units of s^-1^ and they can be more intuitively expressed as:

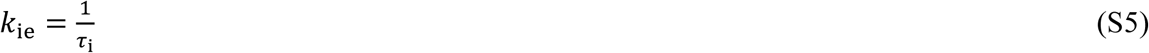

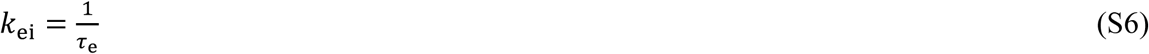

where τ_e_ and τ_i_ are the mean residence times of a substrate molecule in the extra- and intracellular compartments, respectively. Assuming that the volume of an individual cell does not change during the experiment, τ_i_ and *k*_ie_ do not depend on the total number of cells in the sample (determined by the total volume of the sample and the *Ht* value). On the other hand, *k*_ei_ can be calculated from *k*_ie_ as (1):

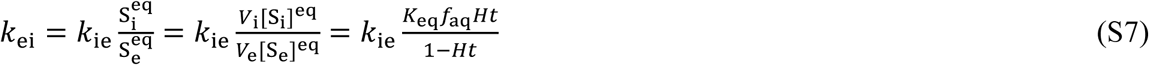

where [S_i_]^eq^ and [S_e_]^eq^ are the equilibrium molar concentrations of the substrate inside and outside the cells, respectively. *k*_eq_ is the equilibrium constant of the membrane transport reaction; for the passive enzyme-mediated diffusion, as in the case of GLUT1, *k*_eq_ = 1 and [S_i_]^eq^ = [S_e_]^eq^. The definitions of *V*_i_, *V*_e_ and *f*_aq_ are as in Eq. S1.

For a carrier-mediated process, the maximum value of the transmembrane flux *J*^eq^ is limited by the capacity and number of the transporter proteins in the cell membrane. Therefore, at a certain high concentration of the substrate, the value of *J*^eq^ will virtually stop increasing upon further additions of the substrate to the medium. It is therefore clear (from Eq. S4) that for a carrier-mediated process, the values of *k*_ie_ and *k*_ei_ depend on the total amount (or concentration) of the substrate in the medium and, in general, they will decrease with increasing substrate concentration.

In the context of the analysis of the 1D-EXSY data, pertinent to this work, the Bloch-McConnell equations relate rate constants *k*_ie_ and *k*_ei_ to the time dependence of the observed changes in the magnetization of the extracellular *M*_e_(*t*) and intracellular *M*_i_(*t*) species during the mixing time of the experiment:

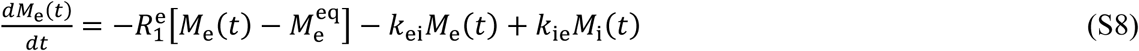

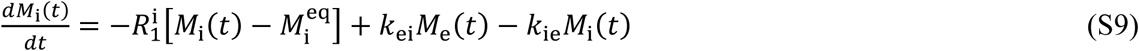

where 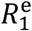 and 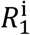 are the longitudinal relaxation-rate constants, characterizing the return of the longitudinal magnetizations to the corresponding thermal-equilibrium values, 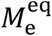 and 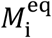.

The appearance of the apparent exchange rate constants *k*_ei_ and *k*_ie_ in Eqs. S8-S9 is justified by the following observations. The magnetizations *M*_i_(*t*) and *M*_e_(*t*) are derived directly from the peak integrals in the NMR spectra, which are proportional to the total mole *amounts* of chemical species in the sample. Since *k*_ei_ and *k*_ie_ relate the transmembrane fluxes of a substrate (in mol s^-1^) and the current (instantaneous) mole amounts of the substrate on either side of the membrane, these are the same rate constants as those used in the Bloch-McConnell equations, which are concerned with the transmembrane fluxes of the two magnetizations (proportional to mole amounts of the two spin populations).

#### Permeability coefficients

The ease with which a substrate is transported across the membrane can also be expressed in terms of the permeability coefficient *P* (3, 4):

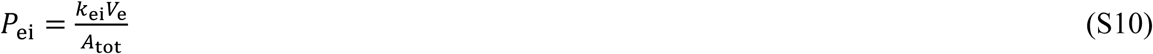

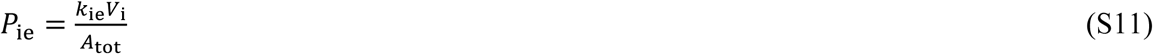

where *P*_ei_ and *P*_ie_ are the permeability coefficients for influx and efflux, respectively; *A*_tot_ is the total surface area of the cell membranes in the sample. As follows from Eq. S7, for GLUT1 at equilibrium: *P*_ei_ = *P*_ie_ (4). However, as in the case of the apparent exchange rate constants, the permeability coefficient is a function of substrate concentration for a carrier-mediated process, and therefore is not ideal for the characterization of transport.

#### Transport rates

The most rigorous way of characterizing carrier-mediated transport kinetics is via determination of reaction rates and their dependence on the substrate concentration, expressed in terms of Michaelis-Menten parameters. In the context of membrane transport, the classical reaction rates are identical to the substrate-transport rates; they are calculated as transmembrane fluxes, normalized to the total aqueous volume of the sample, and measured in units of mol L^-1^ s^-1^ (1):

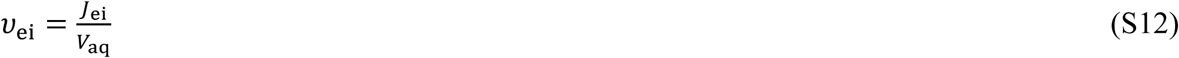

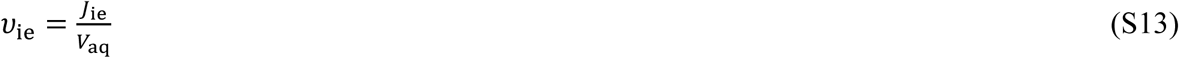

Analogously, for the equilibrium-exchange situation, we obtain the following expression:

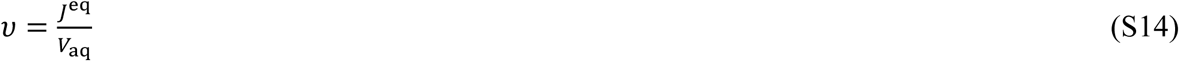

In Eq. S14, omitting the subscript simply indicates the equilibrium-exchange transport rate (*v*), which is the same in both directions.

As alluded to above, when the kinetic process is saturable (enzyme-mediated), the transmembrane rates (as well as fluxes and rate constants) are functions of the substrate concentration. For a true ‘carrier’ transporter, such as GLUT1, the dependence of the transmembrane-transport rates is known to be described well by Michaelis-Menten theory (5). Adaptation of this theory to make it applicable to an equilibrium mixture of two anomeric forms is described next.

### Kinetic theory describing carrier-mediated transmembrane transport

Here we derive the most general form of the equations describing the dependence of carrier-mediated transmembrane-transport rates on substrate concentrations. Next, we assess whether it possible to estimate values of apparent affinities (Michaelis constants *K*_m_) and maximum velocities (*V*_max_) for each of the α- and β-anomers of glucose (and fluorine-labelled analogues of these) in an experiment on RBCs when the system is in thermodynamic equilibrium, but the continuing transmembrane flux is measured by using NMR magnetization-transfer spectroscopy. Finally, we consider the type of kinetic or binding affinity information that could be extracted from the data. In order to set out all the basic terminology, we first consider the kinetics of GLUT1-facilitated transport of *one* substrate across the cell membrane.

### One-substrate GLUT1 kinetics

#### Kinetic model

GLUT1 is known to operate by isomerization: each subunit (that we consider here) of the homotetramer presents its substrate binding site either outside or inside of the cell membrane, in both loaded and unloaded states. This is a feature of a true ‘carrier’ that distinguishes it from a ‘pore’ (such as capnophorin) (5, 6), for which there is only isomerization between the loaded states. For GLUT1, the relative rates of isomerization of the loaded and unloaded states are different, thus the simplest model that describes its transport kinetics is that shown in Fig. S1. There are a total of eight unitary rate constants in the reaction scheme, but there are only seven independent values because of the principle of microscopic reversibility (PMR), which states that the product of rate constants in one direction around a closed reaction loop is equal to the product in the opposite direction (7):

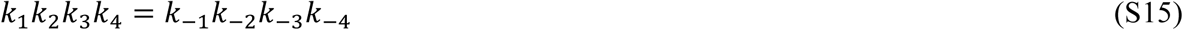

**FIGURE S1.**
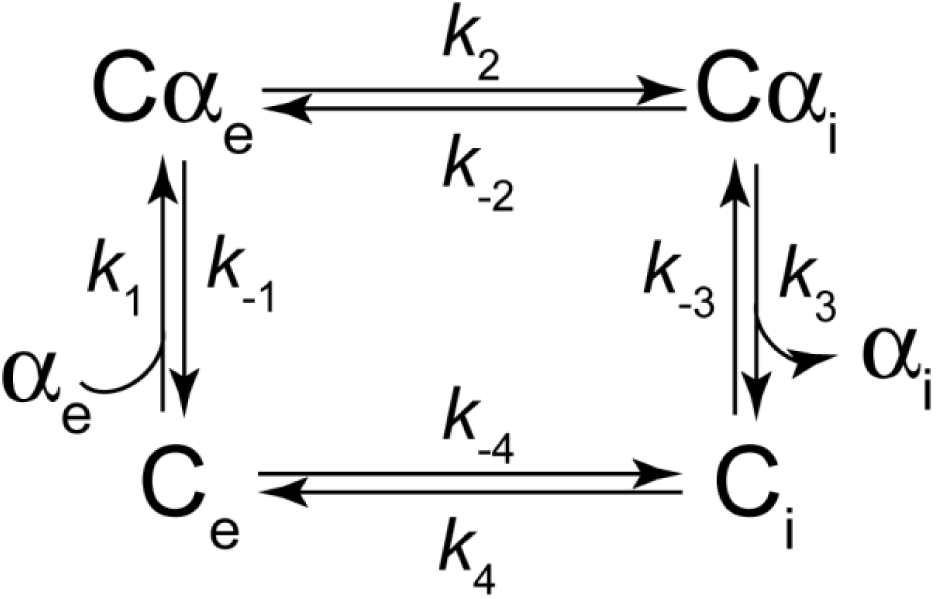
Simplest kinetic model representing transport of the substrate ‘α’ by the carrier protein ‘C’ (GLUT1). Subscripts ‘e’ and ‘i’ specify extra- and intracellular; thus, C_e_ and C_i_ denote the substrate-free carrier, facing either outside (C_e_) or inside the cell membrane (C_i_); Cα_e_ and Cα_i_ are the complexes of the carrier with the substrate, which are facing outside or inside the cell membrane, respectively. The unitary rate constants are denoted by *k*_±i_, i = 1,…,4.

#### Steady-state assumption

In order to derive the equation for the rate of transmembrane exchange, we make the steady-state assumption that the concentrations of the various forms of the carrier remain constant over time (after a very short ‘transient time’ from initiation of a transport experiment):

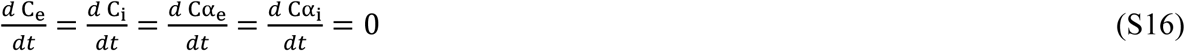

In Eq. S16, C_e_, C_i_, Cα_e_ and Cα_i_ are labels for the four carrier states, and although it is traditional to indicate the concentration of such a state by embracing its label in square brackets, in the interest of reducing clutter in our subsequent expressions, we have left these out. Expressions for the steady-state concentration of each carrier state can be derived by using the method of King and Altman (8), Orsi (9), and Indge and Childs (10) (see examples of derivations for enzymes in (11)). However, our preferred method, and that which appears to make the others redundant (although still relying on drawing a reaction scheme as in (8)), is based on the algorithm of Cornish-Bowden (12), which has been implemented in *Mathematica* (13). The method involves three steps: (a) design the reaction scheme as in Fig. S1; (b) construct a so-called kinetic matrix that is shown below in Fig. S2; and (c) execute the algorithm that we have called ‘rateequationderiver’ in *Mathematica* (13). This generates the requisite expressions for the concentrations of the carrier states, and the corresponding differential equations for reactant flux. In turn, this allows calculation of the transport rate of the substrate, as described below.

**FIGURE S2.**
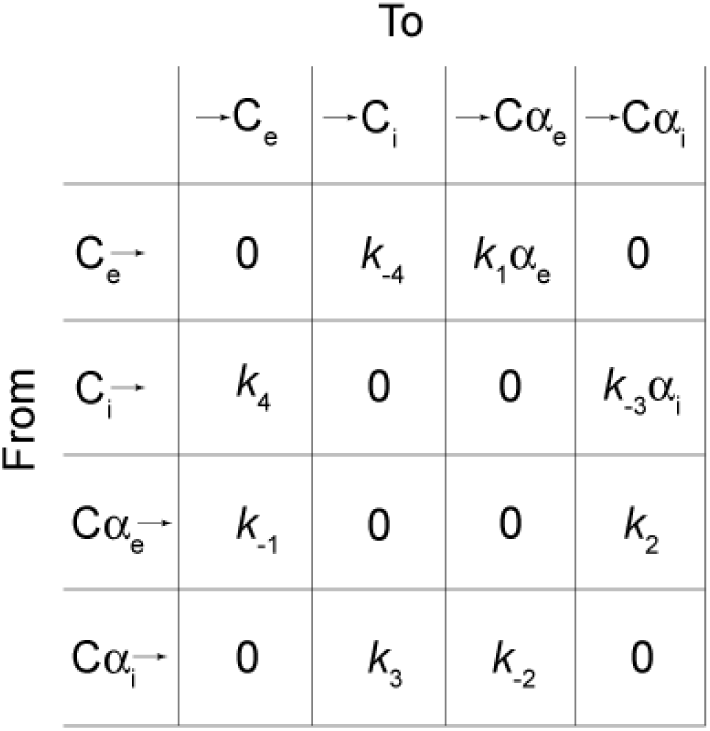
Kinetic matrix corresponding to the reaction scheme of Fig. S1. The matrix is constructed from the list of all states of the carrier, with the elements containing rate constants that characterize the rate of the reaction *from* a state written in the left-hand column, *to* a state in the top row. Hence, the column on the left is labelled ‘From’ and each state has a ‘leaving arrow’ associated with it, while the row at the top is labelled ‘To’ and each state has an arrow directed at it from the left.

The algorithm/function ‘rateequationderiver’ generates the following expressions in terms of the unitary rate constants for the steady-state concentrations of the four carrier species, relative to the total concentration of the carrier:

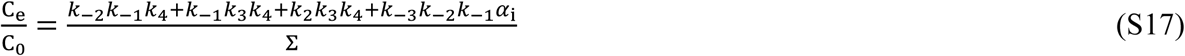

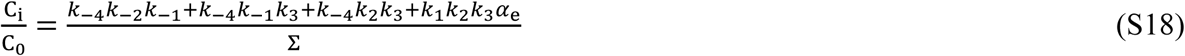

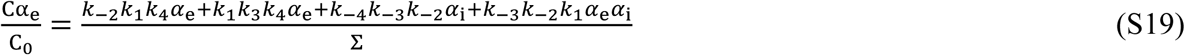

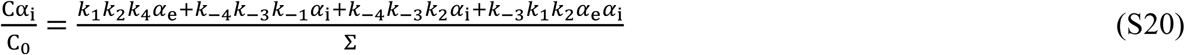

where C_0_ is the total carrier concentration (C_0_ = C_e_ + C_i_ + Cα_e_ + Cα_i_); symbols α_i_ and α_e_ represent concentrations of the substrate outside and inside the cells, respectively; the denominator Σ is the sum of the four numerators of Eqs. S17-S20, and it can be written down as:

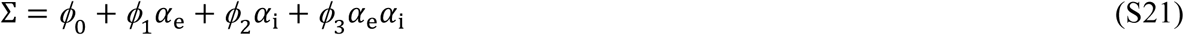

Eq. S21 was obtained by collecting terms that are independent of concentration, then those associated with each solute concentration and the second degree (cross) term. The definitions for the *ø* coefficients are as follows:

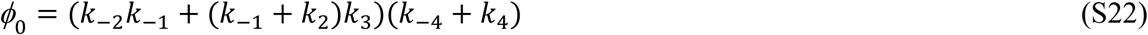

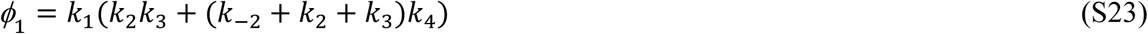

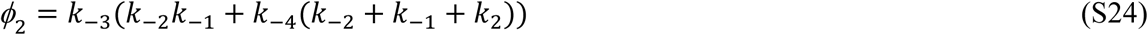

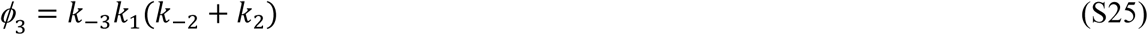

#### Expression for the transmembrane transport rate

To derive the expression for the transport rate of a solute from one side of a membrane to the other via GLUT1, we proceed with the analysis that was first suggested by Britton (14). Consider the flow of substrate from the extra- to the intracellular compartment. The rate at which the substrate forms a complex with the carrier on the extracellular side of the membrane 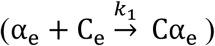 is given by *k*_1_α_e_C_e_. Once the substrate is bound as Cα_e_, it either converts to Cα_i_or dissociates back to α_e_ and 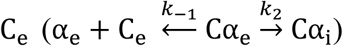. The fraction of the complex Cα_e_ that is converted into Cα_i_ per unit of time (referred to as a transition probability) is a fraction of the total efflux away from 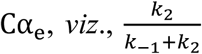. Thus, the overall fraction of substrate flow from α_e_ into Cα_i_ is the product of the two transition probabilities, 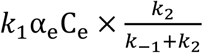. Once the transition αe → Cα_i_ has occurred, the substrate fraction proportional to *k*_3_ is delivered to the other side of the membrane 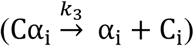, while the amount proportional to *k*_−2_ isomerizes back to 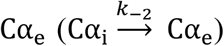; but only the fraction 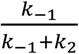 of this goes back to α_e_ and entirely leaves the carrier. Hence, the fraction of substrate bound as Cα_i_ that is delivered to the inside of the cell can be calculated as 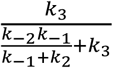. Finally, the expression for the transport rate of the substrate from outside to inside the cells (α_e_ → α_i_) is:

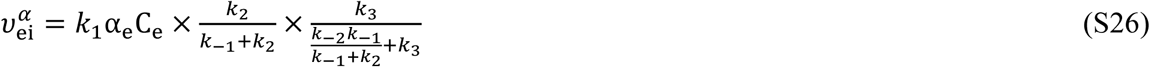

The expression for C_e_ is known from Eq. S17 and upon substituting it into Eq. S26, we obtain:

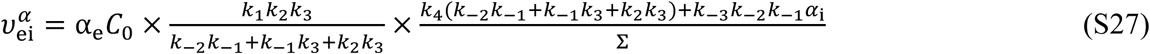

Dividing the numerator and denominator of Eq. S27 by *k*_−3_ *k*_−2_ *k*_−1_ gives:

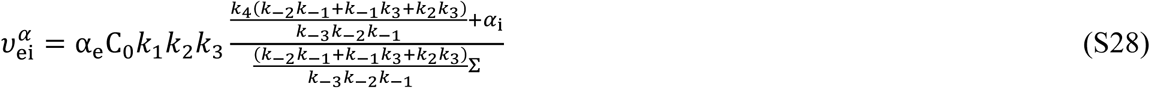

It turns out to be convenient to define the following parameter:

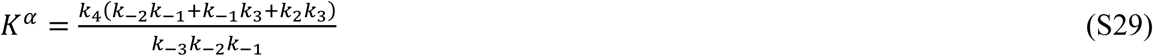

*K^α^* has dimensions of concentration like a Michaelis constant and could be interpreted as an ‘intrinsic affinity constant’ (see below) of the carrier for the substrate *α* (15).

Applying the PMR (7) in the form of Eq. S15 allows simplification of *K^α^* to:

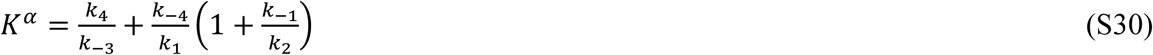

By substituting Eq. S29 into Eq. S28, we obtain the following expression for the transport rate:

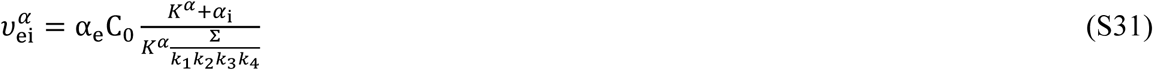

Next, consider the expression.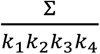. By substituting Eq. S21 into it we obtain:

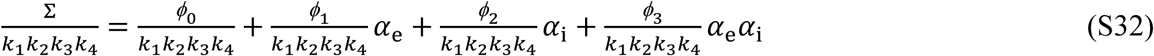

Making the following substitutions brings the terminology in line with that of Stein (5):

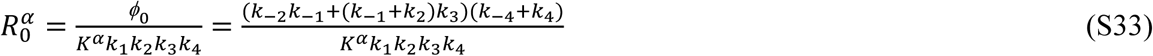

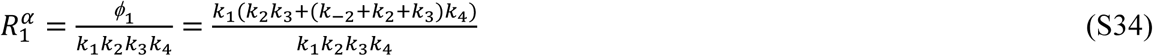

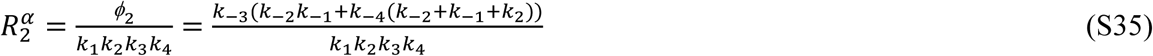

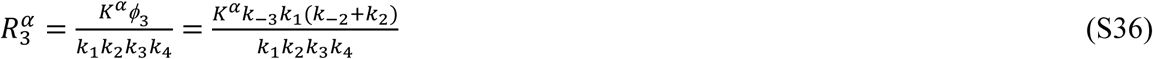

Eqs. S33-S36 are further simplified by using Eq. S15, and the following expressions are obtained:

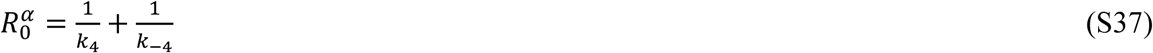

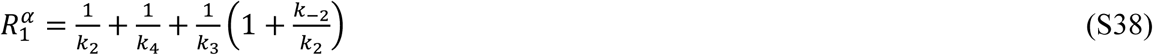

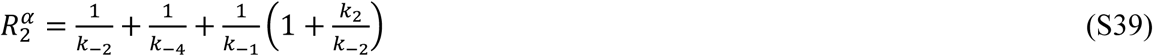

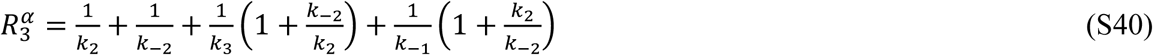

Note that the expression for 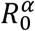 only depends on *k*_4_ and *k*_−4_ (Eq. S37), which are the unitary rate constants that characterize the isomerization of the unloaded carrier, thus its value does not depend on the nature of the substrate, so we can drop the superscript α from it. The physical meaning of *R*_0_ is the average time (also called the life time) that it takes for the carrier to jump from one unloaded state to the other and then back (in the absence of any substrate).

From the expressions in Eqs. S37-S40, it can be readily seen that:

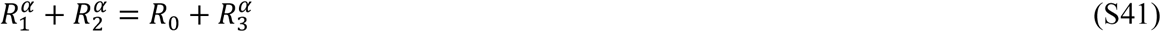

Substituting Eqs. S33-S36 into Eq. S32 gives:

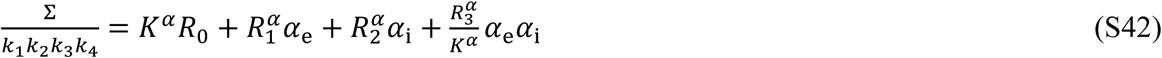

Finally, substituting Eq. S42 into Eq. S31 yields the following expression for the transport rate:

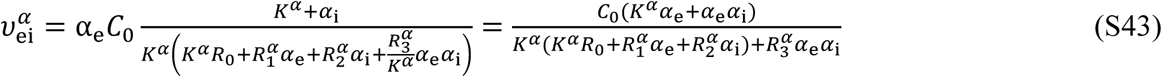

Thus, Eq. S43 is the ‘master equation’ describing the forward (from extracellular ‘e’ to intracellular ‘i’) transport rate, 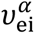, in its most general form. Eq. S43 is fully analogous to Equation (4.2) of Stein (5) Definitions of parameters 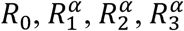 and *K*^*α*^ in terms of the unitary rate constants of Figure 1 (Eq. S30 and Eqs. S37-S40), are also completely equivalent to those in Table 4.1 of Stein (5). By recognizing the symmetry in the reaction scheme, the equation for the reverse rate of transport, 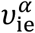, is obtained by swapping the subscripts ‘i’ and ‘e’:

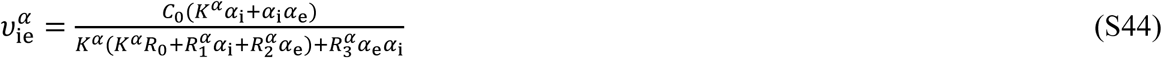

#### Specific case of equilibrium-exchange kinetics

In the present paper, we focus on the specific case of equilibrium-exchange transport. Because of the very rapid transmembrane equilibration of glucose (and its fluorine-labelled analogues) by facilitated diffusion via GLUT1, the concentrations of the anomers become identical inside and outside of the cells. This is the basic *equilibrium-exchange* condition (5, 6):

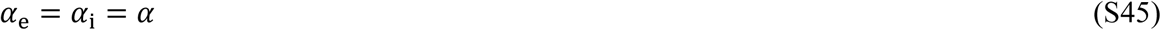

Upon substituting Eq. S45 into the master equation (Eq. S43), we obtain the expression for the transport rate under equilibrium-exchange conditions:

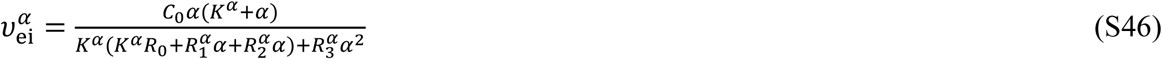

Note that right-hand expression of Eq. S46 would be the same for the transport rate in the opposite direction (if we substitute Eq. S45 into Eq. S44 instead), thus, as in Eq. S14, we can drop the subscript and simply define the equilibrium-exchange rate of transport as *v^α^*:

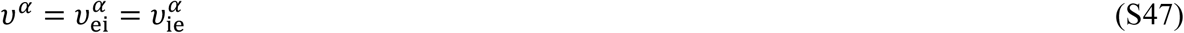

Eq. S46 can be further simplified by using the equality of Eq. S41:

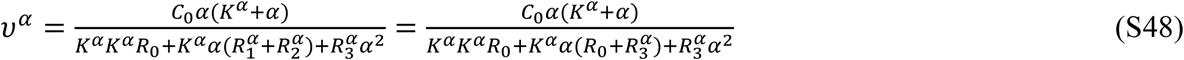

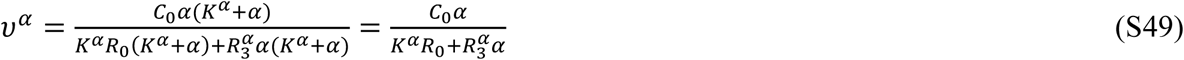

In order to cast Eq. S49 into the form of the classical Michaelis-Menten equation, we divide the numerator and denominator by 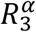:

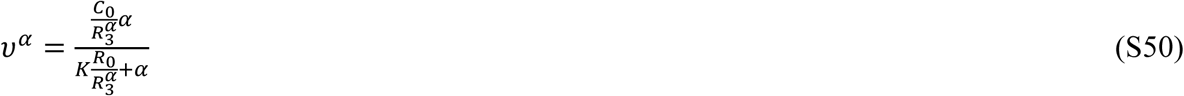

Thus, in the case of equilibrium-exchange transport, the carrier-mediated transport is fully described by Michaelis-Menten kinetics and the parameters (relevant to the equilibrium-exchange case) can be expressed as (in full analogy with the Table 4.2 of Stein (5)):

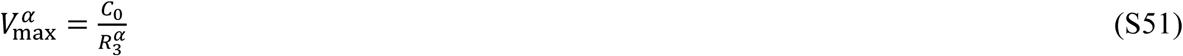

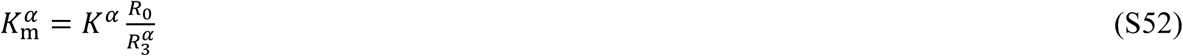

Thus, as for standard Michaelis-Menten enzyme kinetics, it would be possible to estimate values of apparent affinities (*K*_m_) and maximum velocities (*V*_max_) for *one* GLUT1 substrate in an experiment, when the system is in thermodynamic equilibrium. The experiment would involve measuring the transmembrane-transport rate in either direction (the rates are equal in both directions) for a range of substrate concentrations and fitting Eq. S50 to the data that reveal the dependence of the transport rate on the substrate concentration.

Having estimated the values of the parameters 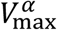 and 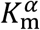 we can, in principle, estimate the value of 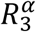, which is the inverse of the turnover number, 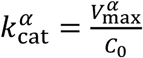. The reciprocal of this signifies the average time for the carrier to transport one molecule of substrate *α*, under saturated conditions. Aside: analogously, it can be shown that 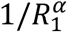 is the average time for the carrier to transport one molecule of substrate from outside to inside the cell, when all of the substrate is located outside; 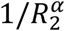 is the average time for the carrier to transport one molecule of substrate from inside to outside the cells, when all of the substrate is located inside.

However, in the experiments reported in the present work, there is a further restriction imposed on the analysis by the presence of the second competing substrate. In fact, the two anomeric species are always present together as a racemic mixture in a fixed ratio, which for native glucose in water is α-anomer:β-anomer = 36:64. In the ensuing kinetic analysis, we consider the kinetics of the two competing substrates, and the anomerization rate is assumed to be negligible during the actual membrane passage of a given anomer (see Results for evidence of this assumption).

### Two-substrate GLUT1 kinetics

#### Kinetic model

Now consider the kinetics of transport of *two* competing substrates *α* and *β* across the cell membrane. With two anomers present, there are now a total of six states of the carrier (four loaded and two unloaded). The kinetic scheme for such transport is shown in Fig. S3 (in analogy with Fig. 4.11(a) of Stein (5)):

**FIGURE S3.**
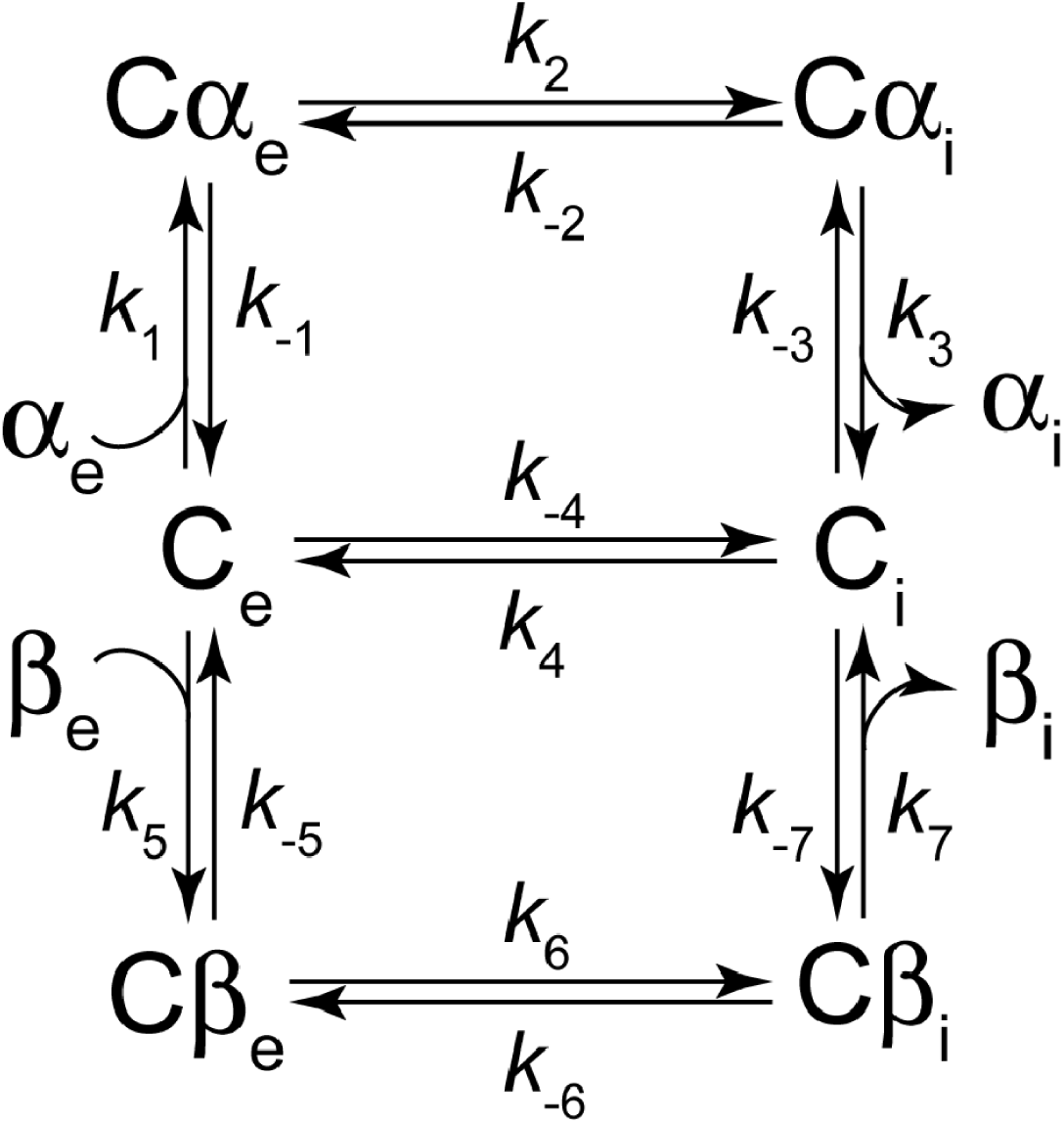
Two-substrate kinetic scheme representing a simple model of the transport mechanism of GLUT1. α and β denote the respective anomers of glucose, ‘e’ and ‘i’ specify extra- and intracellular. C is the carrier protein (GLUT1) and, when juxtaposed to α or β, the symbol denotes the respective protein-substrate complex. The unitary rate constants are denoted by *k*_±i_, i = 1,…,7 giving a total of 14 in the scheme.

As in the one-substrate scheme (Fig. S1), applying the PMR we obtain the following two equalities:

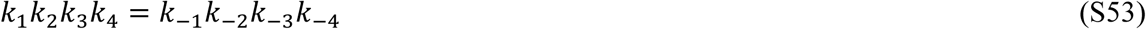

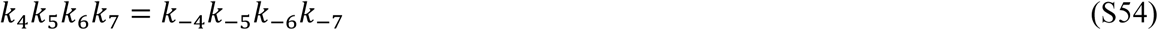

Additionally, by combining Eqs. S53-S54, a similar constraint for the larger kinetic loop is identified:

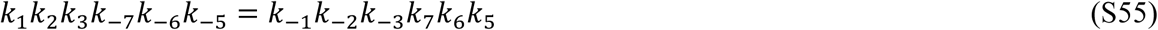

As before, we compose the corresponding kinetic matrix, shown in Fig. S4.

**FIGURE S4.**
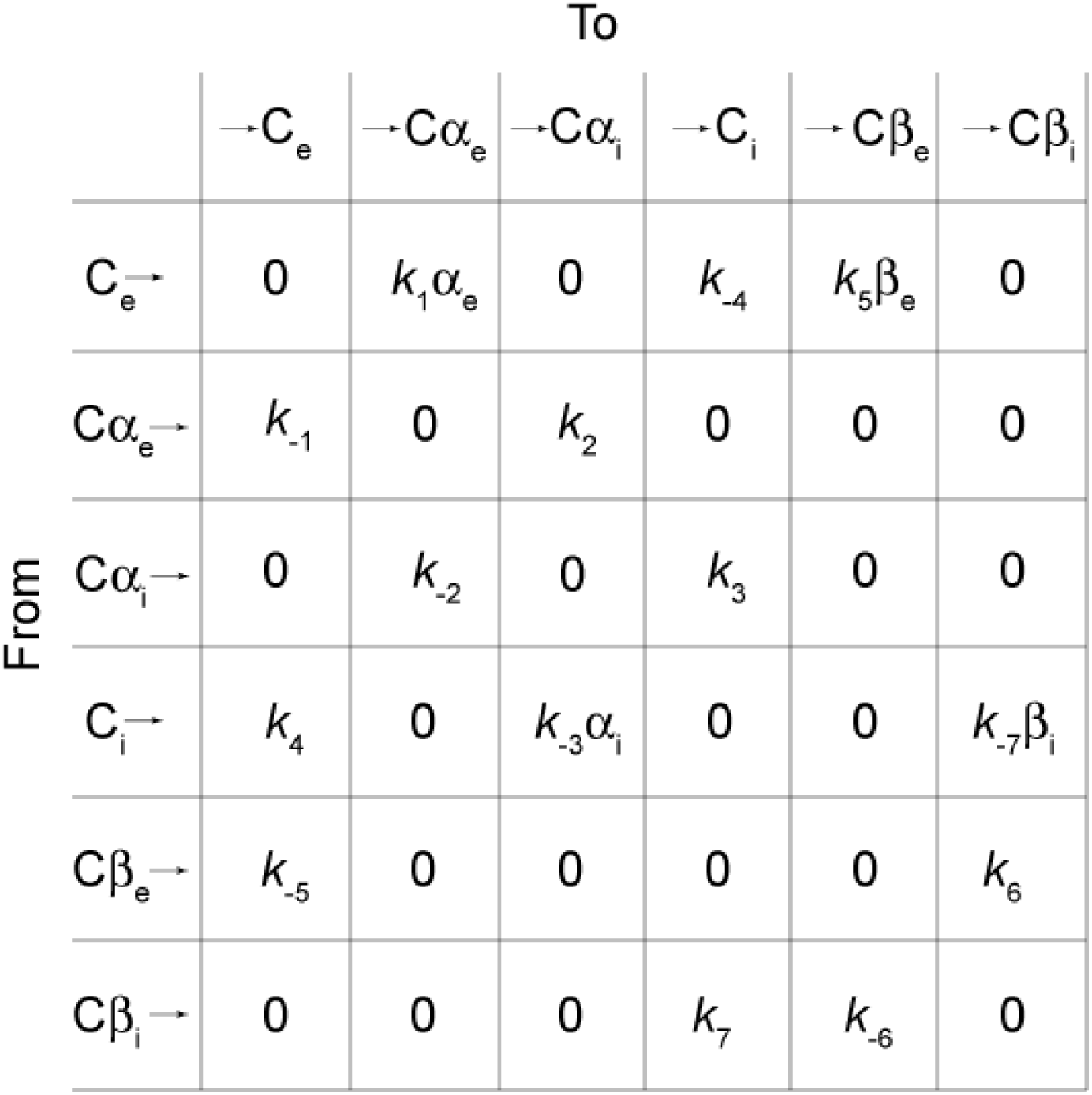
Kinetic matrix corresponding to the reaction scheme of Fig. S3. It is constructed from the list of all states of the carrier, with the elements containing rate constants that characterize the rate of the reaction *from* a state in the left-hand column, *to* a state in the top row. Hence, the column on the left is labelled ‘From’ and each state has a ‘leaving arrow’ associated with it, while the row at the top is labelled ‘To’ and each state has an arrow directed at it from the left.

The logic guiding the derivation of the expression for the transport rate remains the same as in the one-substrate scheme. Thus, by analogy with Eq. S26, we obtain the expression for the transport rate of each anomer from outside to inside the cells:

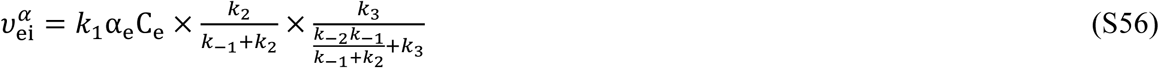

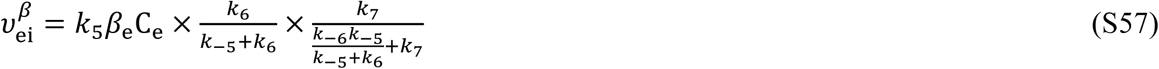

Using the kinetic matrix of Fig. S4, ‘rateequationderiver’ (13) in *Mathematica* delivers the following expression for the required steady-state concentration of unloaded carrier facing outside (C_e_):

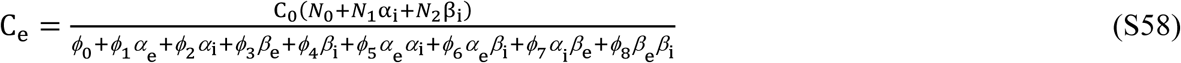

In Eq. S58, the following substitutions were made:

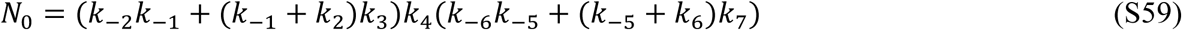

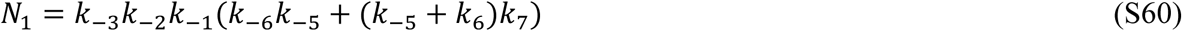

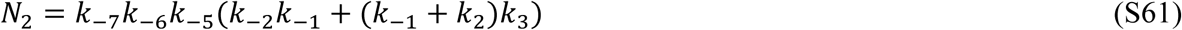

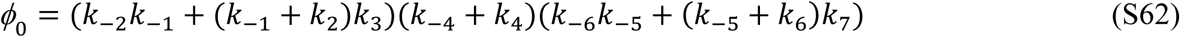

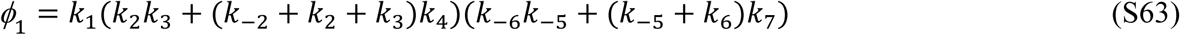

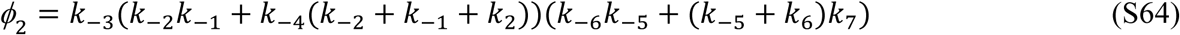

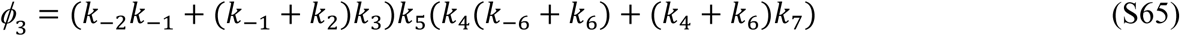

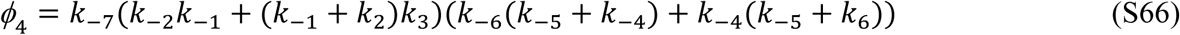

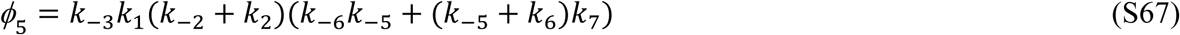

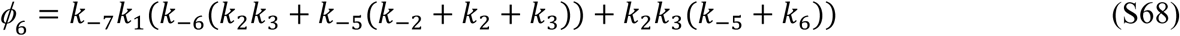

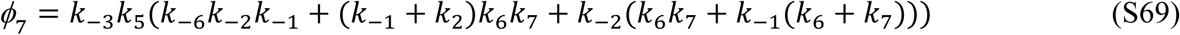

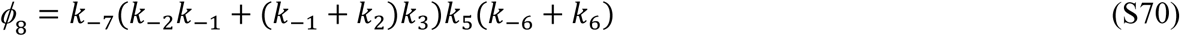

Next, consider the equation for the transport of just one substrate (*e.g*., *α*); the following analysis is entirely analogous for the other competing anomer. Substituting Eq. S58 into Eq. S56 yields:

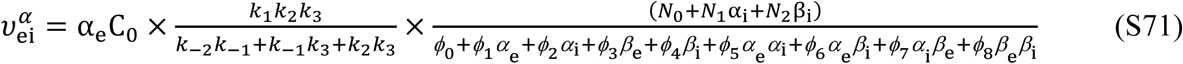

Using the definition of *K^α^* (Eq. S29), we define *K^β^* analogously by referring to Fig. S3:

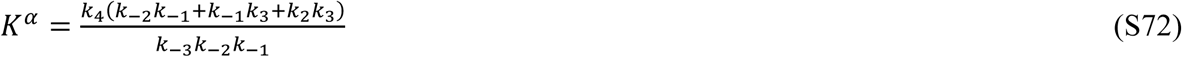

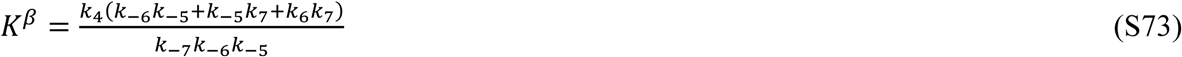

It turns out to be convenient to introduce the following ‘de-dimensionalizing’ (also called scaling) definitions:

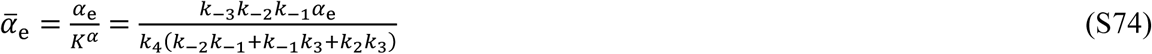

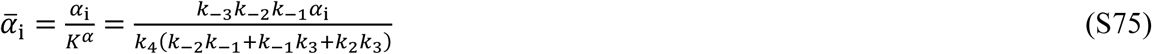

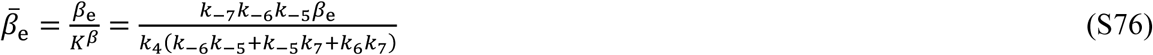

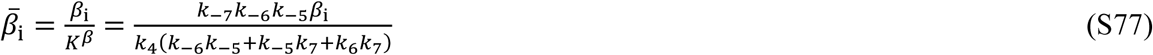

Thus, using, Eqs. S72-S77, Eq. S71 is expressed as:

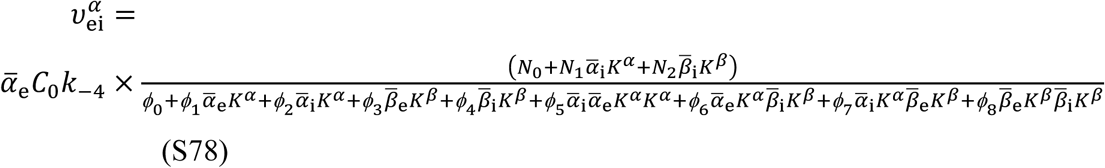

Next, dividing the numerator and denominator by ***N***_0_ and *k*_−4_ yields:

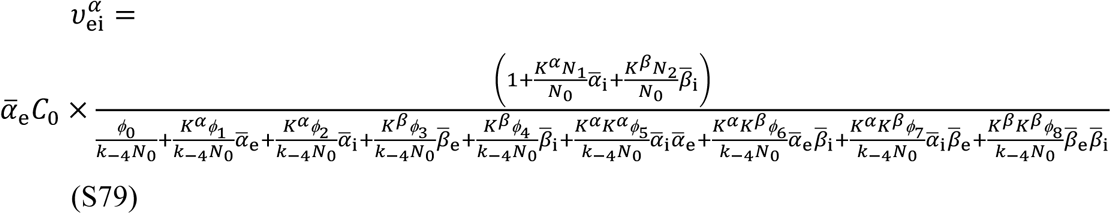

Now consider each of the coefficients (there are some striking mathematical simplifications here) in the numerator and denominator of Eq. S79. In the process, introduce the new definitions:

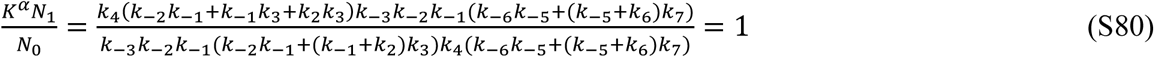

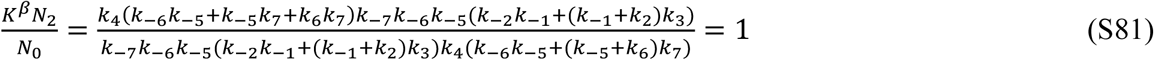

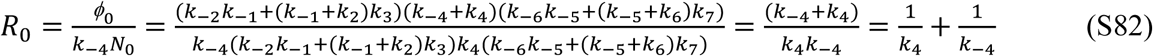

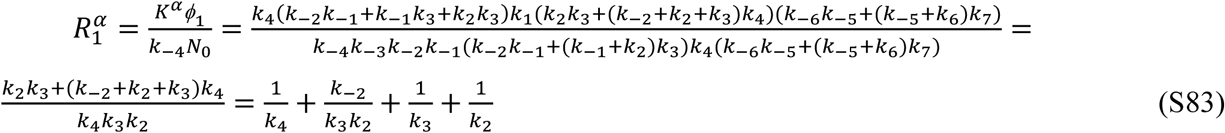

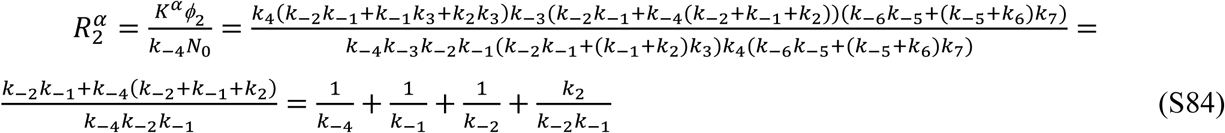

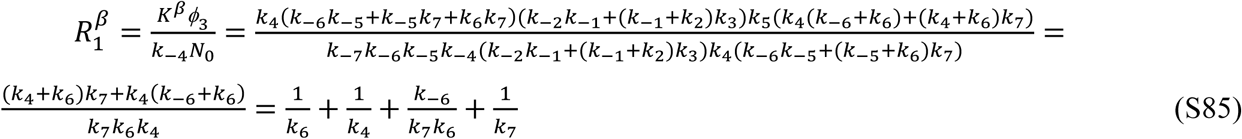

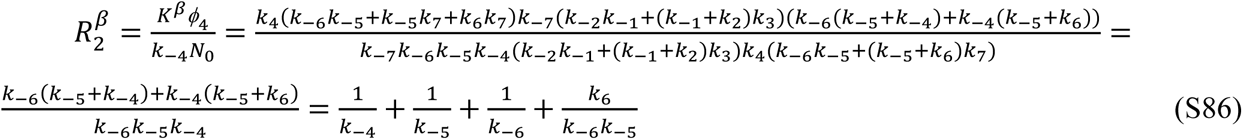

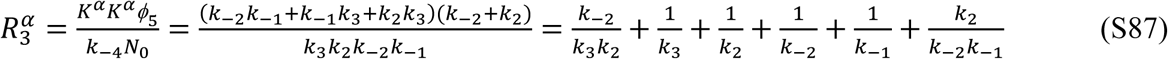

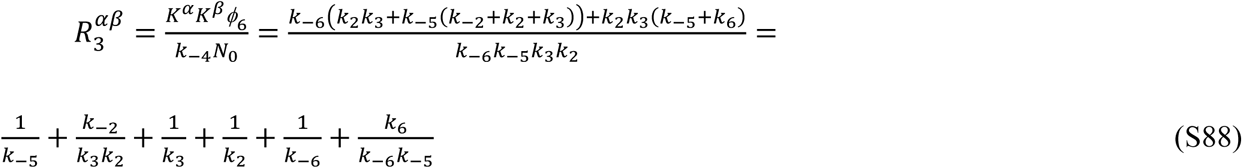

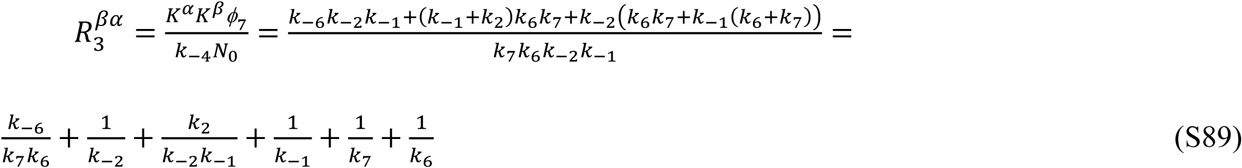

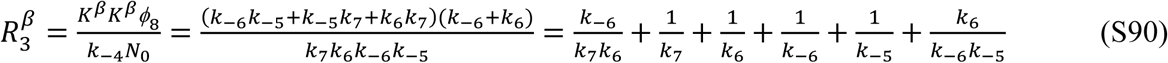

Importantly, note that the expressions for 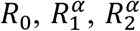, and 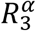 are exactly the same as in the case of the one-substrate transport scheme (Eqs. S37-S40). Moreover, the definitions of 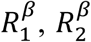 and 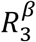 are entirely analogous to 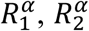 and 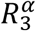 (according to Fig. S3), while parameters 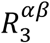 and 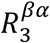 are only relevant when the two competing substrates are simultaneously present in the system; they are referred to as the ‘heteroexchange’ terms (15). According to the definitions of Eqs. S82-S90, we obtain the following equalities:

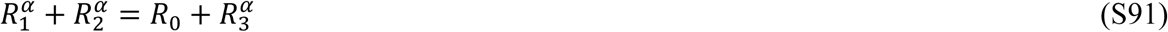

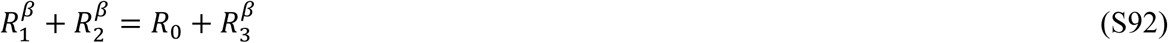

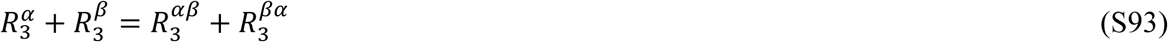

Upon substituting Eqs. S80-S90 into Eq. S79 we arrive at the most general form of the equation for the rate of transport of one substrate (*α*) by the carrier from one side of the membrane (extracellular one) to the other side (intracellular one), in the presence of the competing substrate *β*:

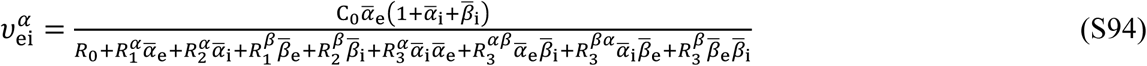

Note that Eq. S94 is completely analogous to Equation (4.23) of Stein (5). The definitions of symbols *K* and *R* are also analogous to the definitions of Table 4.16 of Stein (5), however the *R* symbol subscripts are now different and use the following mappings 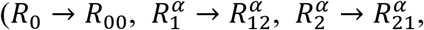 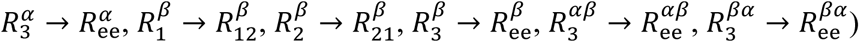.

#### Specific case of equilibrium-exchange kinetics

Now consider the specific case of equilibrium exchange transport. GLUT1 operates by ‘facilitated diffusion’ implying that the transport is energy-independent. Consequently, at equilibrium the concentration of the α-anomer is equal inside and outside the cells and this also applies to the β-anomer. Thus, the concentration of the two substrates are the same on both sides of the membrane:

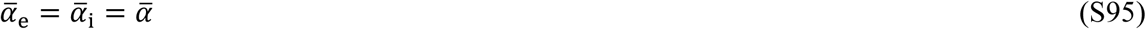

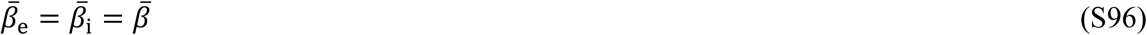

Therefore, we substitute Eqs. S95-S96 into Eq. S94 to obtain:

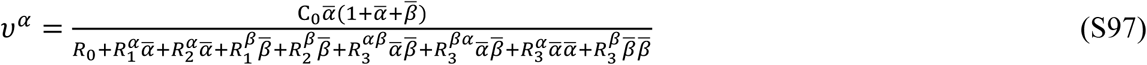

As for the one-substrate case, the transport rates in each direction are the same, thus to reduce complexity we use the symbol *v^α^* rather than 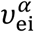. By analogy, we could have also written the equation for the β-anomer, with the following analysis being completely identical. Eq. S97 can be further simplified to:

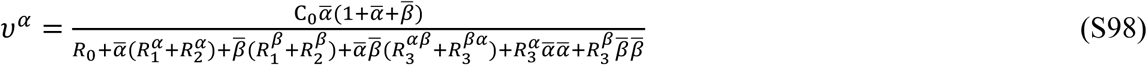

Introducing the equalities of Eqs. S91-S93 into Eq. S98 yields:

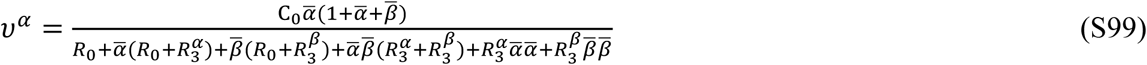

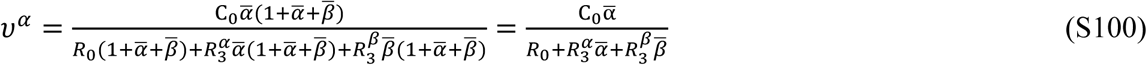

Thus, the following equation for the equilibrium-exchange transport rate of *α* is obtained by rescaling back to α and β using Eqs. S74-S77:

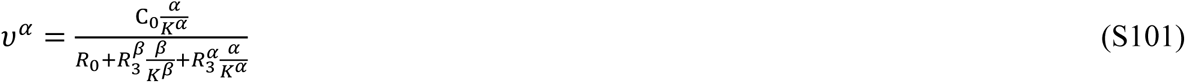

This equation is cast into the form of the Michaelis-Menten expression by multiplying the numerator and denominator by 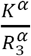:

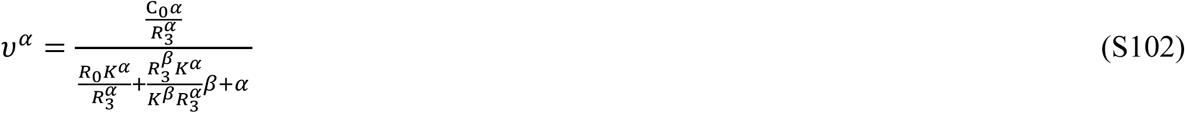

Analogously, for the transport rate of *β* the following expression applies:

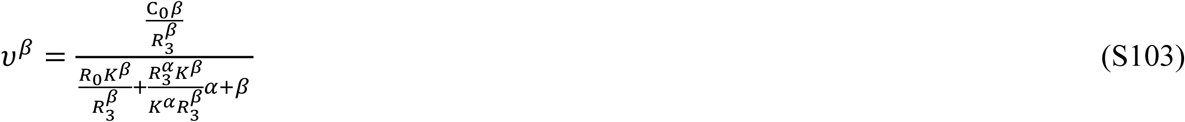

Return to Eqs. S51-S52 to be reminded that the one-substrate equilibrium-exchange Michaelis-Menten parameters are:

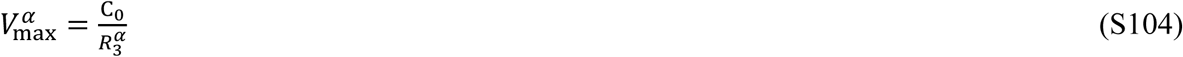

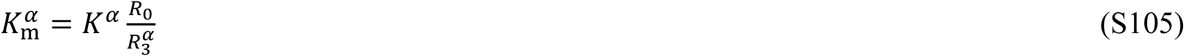

By analogy, for the second substrate, the kinetic parameters are:

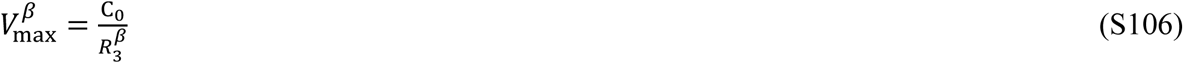

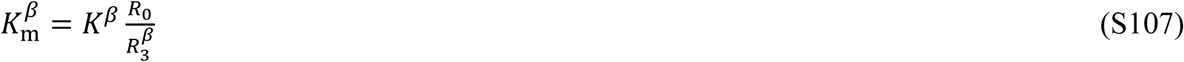

Substituting Eqs. S104-S107 into Eqs. S102-S103, yields the following new ‘master equations’:

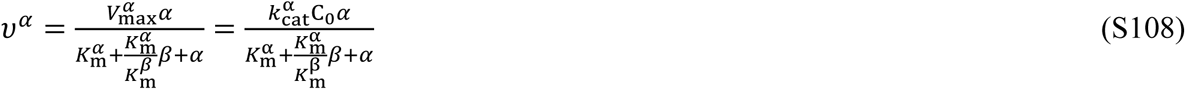

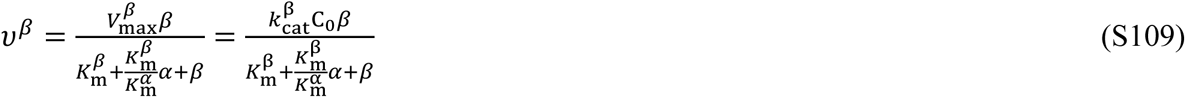

where 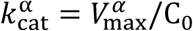 and 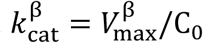 are the turnover numbers of GLUT1 for the two anomers (16).

Note that the values of *V*_max_ and *k*_m_ in Eqs. S104-S109 are specific for the *equilibrium-exchange* condition. Equilibrium-exchange kinetics is closer to the physiological situation than are *zero-cis*, *zero-trans*, *infinite-cis* and *infinite-trans* experiments, where kinetics are measured before transmembrane equilibrium is reached (5). In non-equilibrium kinetics, the expressions for the Michaelis-Menten parameters are different for efflux and influx – they can be derived in analogy with the approach used here and elsewhere (17, 18). However, under equilibrium-exchange, pertinent to the present work, the Michaelis-Menten expressions are the same for the two directions of transport (19).

Eqs. S108-S109 are quite simple and, in fact, can be recognized as classical Michaelis-Menten equations for two competing substrates (3). Under normal conditions (minimal or zero mutarotase activity, or lack of spontaneous mutarotation), the rate of interconversion is slow on the NMR timescale (see Figs. 5-7 in the main text), so at equilibrium, the ratio of the concentrations of the two anomers in each compartment is constant and can be defined by parameter *a*, or, alternatively, by the equilibrium constant of mutarotation, *k*_mut_:

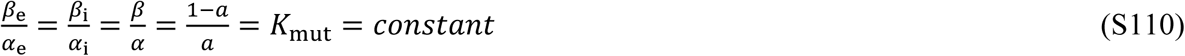

Substitutions [α] *= aS* and [β] = (1 − *a*)*S* (where *S* = α + β is the total substrate concentration) into Eqs. S108-S109, followed by normalization of the transport rates by the total carrier and substrate concentrations, gives:

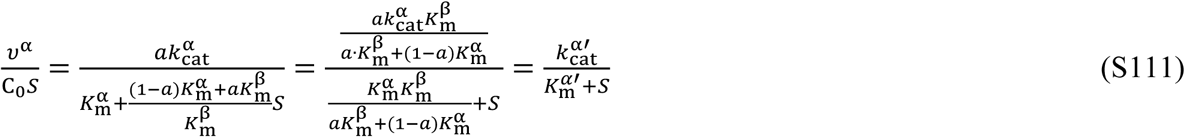

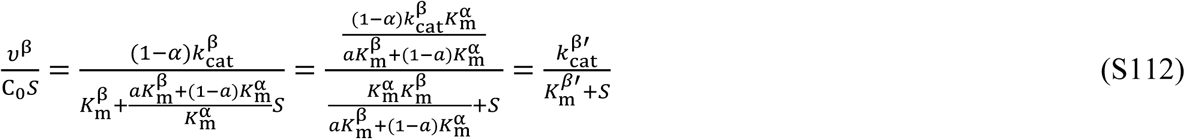

In Eqs. S111-112, the following definitions are made in order to obtain the ‘apparent’ Michaelis-Menten equations:

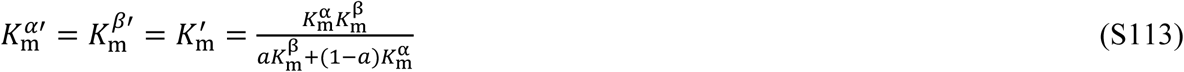

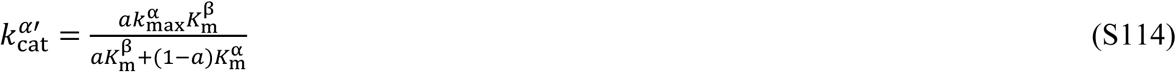

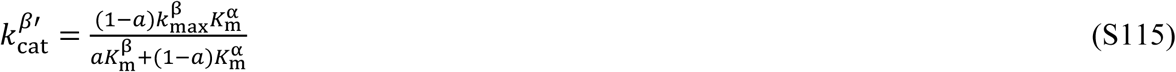

Eqs. S111-S112 have the form of the classical Michaelis-Menten equations. Thus, the apparent Michaelis-Menten parameters (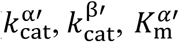 and 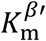) could be determined by fitting the model of Eqs. S111-S112 to the experimental data (dependence of the normalized transport rates on the substrate concentration in the equilibrium-exchange experiment). Interestingly, from Eqs. S111-S115, it follows that:

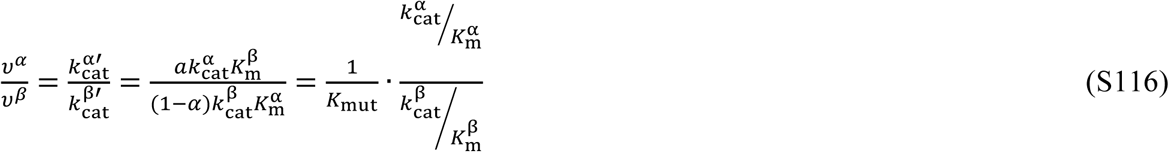

Thus, the ratio of the rates of transport of the competing anomers is constant, irrespective of the total concentration of the substrate *S*, and defined by three constants: *k*_mut_ and the two specificity constants *k*_cat_ /*k*_m_ of the carrier for each of the competing anomers. Substituting Eqs. S104-S107 into Eq. S116 yields:

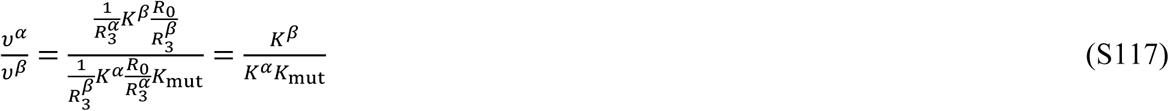

Thus, constants *K^α^* and *K^β^* define the ‘intrinsic affinity constant’ of the carrier towards the two anomeric substrates (15).

Experimentally, we are able to measure the rate of exchange of both the α- or β-anomers of glucose-derivatives and record the concentration-dependence of these rates. Then, if the data are graphed according to the classical steady-state enzyme-kinetic analysis, it will allow estimation of the values of the apparent Michaelis-Menten parameters 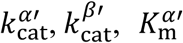 and 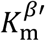 using Eqs. S111-S112. However, while expression for 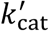 differ for the two anomers (Eqs. S114-S115), those for 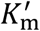 are formally the same (Eq. S113). In other words, it is not possible to estimate the relative affinities of the GLUT1 transporter for both anomers using this approach. On the other hand, consider the expressions since the 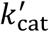 values can, in principle, differ, we can calculate the following expressions: 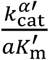 and 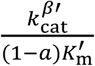. By using Eqs. S113-S115, these can be shown to correspond to the following identities:

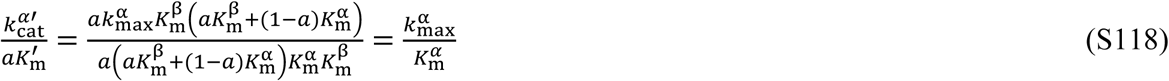

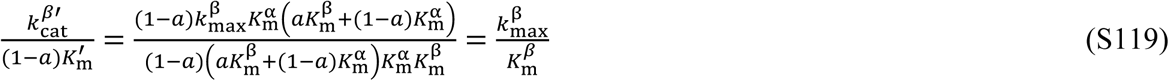

Thus, the specificity constant of each of the two anomers, 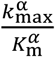 and 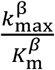 emerge from the data analysis. These values correspond to the normalized transmembrane transport rates of the sugar in the limit of low substrate concentration (*e.g*., the situation when there is an unlimited number of carriers available for transport). In terms of Stein’s nomenclature (5), these values are equal to:

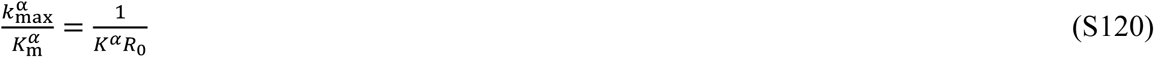

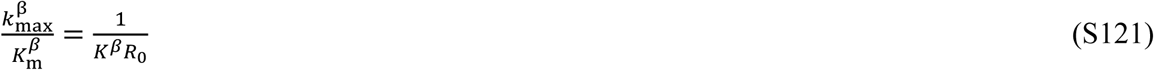

In each of these expressions (Eqs. S120-S121), there is only one parameter (*K^α^* or *K^β^*) that depends on the physico-chemical nature of the substrate, since *R*_0_ is determined by the values of *k*_4_ and *k*_−4_, which do not have any relevance to the nature of the substrate. Thus, these specificity constants are useful quantities for characterizing the overall efficiency of the transport of particular sugar anomers by the carrier protein (GLUT1); they are commonly used by enzyme kineticists when characterizing the catalytic action of enzymes.

To summarize, we have arrived at simple expressions for the rates of transport of two competing GLUT1 substrates under equilibrium-exchange conditions, *v*^*α*^ and *v*^*β*^ (Eqs. S108-S109). The rate equations were further developed for the case when the ratio of the two competing substrates remains constant, as in the case of the two sugar anomers. In this case, rearrangement of the kinetic terms (Eqs. S111-S112) suggests determination of the values of 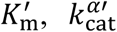 and 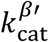 by measuring the transport rates of the two anomers for a range of substrate concentrations and fitting Eqs. S111-S112 to the experimental data. However, it is not possible to obtain estimates of the individual values of *k*_m_ and *k*_cat_ (or *V*_max_) for each anomer without introducing additional assumptions, because anomerization prevents either working with individual anomeric species or varying the anomer ratio. This problem could be resolved in the specific case of monosaccharides which are fluorinated at the anomeric carbon atom (20). Nevertheless, it is significant that, in the general case, Eqs. S118-S119 allow estimation of the individual specificity constants of GLUT1 for the two anomers by using only the experiments in which they are in a constant ratio and are simultaneously interacting with the carrier under equilibrium-exchange conditions.

### Conclusions

As has been shown here, in answer to the opening question, it is not possible to extract individual values of *k*_cat_, *V*_max_ and *k*_m_ from magnetization-transfer NMR experiments (under equilibrium-exchange conditions), with two competing substrates that are always present in solution in a constant ratio. However, when comparing different substituted (F-labelled) glucoses, relative rates could differ for each of them, and since the value of *R*_0_ is independent of the solute being transported, the estimated values of the specificity constants 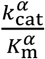 and 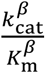 will reflect the relative affinities for GLUT1 between the two anomers, and, in general, between different F-substituted glucoses. In other words, NMR-based magnetization-transfer analysis is a valid method for comparing affinities of F-labelled glucoses by GLUT1, provided the experiments are conducted with ranges of concentrations of the F-labelled glucoses. This is the approach that we have adopted in the present paper.

## REFERENCES

1. Nelson, D. L., and M. M. Cox. 2008. Lehninger Principles of Biochemistry. W.H. Freeman and Company, New York.

2. Carruthers, A. 1990. Facilitated diffusion of glucose. Physiol. Rev. 70:1135–1176.

3. Thorens, B., and M. Mueckler. 2010. Glucose transporters in the 21st Century. Am. J. Physiol.: Endocrinol. Metab. 298:E141–E145.

4. Mueckler, M., and B. Thorens. 2013. The SLC2 (GLUT) family of membrane transporters. Mol. Aspects Med. 34:121–138.

5. Gorga, F. R., and G. E. Lienhard. 1982. Changes in the intrinsic fluorescence of the human erythrocyte monosaccharide transporter upon ligand binding. Biochemistry 21:1905–1908.

6. Macheda, M. L., S. Rogers, and J. D. Best. 2005. Molecular and cellular regulation of glucose transporter (GLUT) proteins in cancer. J. Cell. Physiol. 202:654–662.

7. Ganapathy, V., M. Thangaraju, and P. D. Prasad. 2009. Nutrient transporters in cancer: relevance to Warburg hypothesis and beyond. Pharmacol. Ther. 121:29–40.

8. Amann, T., and C. Hellerbrand. 2009. GLUT1 as a therapeutic target in hepatocellular carcinoma. Expert Opin. Ther. Targets 13:1411–1427.

9. Adekola, K., S. T. Rosen, and M. Shanmugam. 2012. Glucose transporters in cancer metabolism. Curr. Opin. Oncol. 24:650–654.

10. Ung, P. M.-U., W. Song, L. Cheng, X. Zhao, H. Hu, L. Chen, and A. Schlessinger. 2016. Inhibitor Discovery for the Human GLUT1 from Homology Modeling and Virtual Screening. ACS Chem. Biol. 11:1908–1916.

11. Granchi, C., S. Fortunato, and F. Minutolo. 2016. Anticancer agents interacting with membrane glucose transporters. MedChemComm 7:1716–1729.

12. Calvaresi, E. C., and P. J. Hergenrother. 2013. Glucose conjugation for the specific targeting and treatment of cancer. Chem. Sci. 4:2319–2333.

13. Deng, D., C. Xu, P. Sun, J. Wu, C. Yan, M. Hu, and N. Yan. 2014. Crystal structure of the human glucose transporter GLUT1. Nature 510:121–125.

14. Deng, D., P. Sun, C. Yan, M. Ke, X. Jiang, L. Xiong, W. Ren, K. Hirata, M. Yamamoto, and S. Fan. 2015. Molecular basis of ligand recognition and transport by glucose transporters. Nature 526:391–396.

15. Champagne, P. A., J. Desroches, and J.-F. Paquin. 2015. Organic fluorine as a hydrogen-bond acceptor: recent examples and applications. Synthesis 47:306–322.

16. Street, I. P., C. R. Armstrong, and S. G. Withers. 1986. Hydrogen bonding and specificity. Fluorodeoxy sugars as probes of hydrogen bonding in the glycogen phosphorylase-glucose complex. Biochemistry 25:6021–6027.

17. Withers, S. G., I. P. Street, and M. D. Percival. 1988. Fluorinated carbohydrates as probes of enzyme specificity and mechanism. In Fluorinated Carbohydrates. N. F. Taylor, editor. American Chemical Society, Washington, pp. 59–77.

18. Glaudemans, C. P. 1991. Mapping of subsites of monoclonal, anti-carbohydrate antibodies using deoxy and deoxyfluoro sugars. Chem. Rev. 91:25–33.

19. Potts, J. R., A. M. Hounslow, and P. W. Kuchel. 1990. Exchange of fluorinated glucose across the red-cell membrane measured by 19F-NMR magnetization transfer. Biochem. J. 266:925–928.

20. Potts, J. R., and P. W. Kuchel. 1992. Anomeric preference of fluoroglucose exchange across human red-cell membranes: 19F-NMR studies. Biochem. J. 281:753–759.

21. O’Connell, T. M., S. A. Gabel, and R. E. London. 1994. Anomeric dependence of fluorodeoxyglucose transport in human erythrocytes. Biochemistry 33:10985–10992.

22. Dickinson, E., J. R. P. Arnold, and J. Fisher. 2017. Determination of glucose exchange rates and permeability of erythrocyte membrane in preeclampsia and subsequent oxidative stress-related protein damage using dynamic-19F-NMR. J. Biomol. NMR 67:145–156.

23. Dacie, J. V., and S. M. Lewis. 1975. Practical Haematology. Churchill Livingstone, Edinburgh.

24. Shaka, A., J. Keeler, T. Frenkiel, and R. Freeman. 1983. An improved sequence for broadband decoupling: WALTZ-16. J. Magn. Reson. 52:335–338.

25. Claridge, T. D. 2009. High-resolution NMR Techniques in Organic Chemistry. Elsevier, Amsterdam.

26. Wolfram, S. 2003. The Mathematica Book. Wolfram Media, Champaign.

27. Brown, F. F., I. D. Campbell, P. W. Kuchel, and D. L. Rabenstein. 1977. Human erythrocyte metabolism studies by 1H spin echo NMR. FEBS Lett. 82:12–16.

28. Kim, H. W., P. Rossi, R. K. Shoemaker, and S. G. DiMagno. 1998. Structure and transport properties of a novel, heavily fluorinated carbohydrate analogue. J. Am. Chem. Soc. 120:9082–9083.

29. Bresciani, S., T. Lebl, A. M. Z. Slawin, and D. O’Hagan. 2010. Fluorosugars: synthesis of the 2,3,4-trideoxy-2,3,4-trifluoro hexose analogues of D-glucose and D-altrose and assessment of their erythrocyte transmembrane transport. Chem. Commun. 46:5434–5436.

30. Kuchel, P. W. 1981. Nuclear magnetic resonance of biological samples. Crit. Rev. Anal. Chem. 12:155–231.

31. Bloch, R. 1973. Inhibition of glucose transport in the human erythrocyte by cytochalasin B. Biochemistry 12:4799–4801.

32. Robinson, G., P. W. Kuchel, B. E. Chapman, D. M. Doddrell, and M. G. Irving. 1985. A simple procedure for selective inversion of NMR resonances for spin transfer enzyme kinetic measurements. J. Magn. Reson. 63:314–319.

33. McConnell, H. M. 1958. Reaction rates by nuclear magnetic resonance. J. Chem. Phys. 28:430–431.

34. Shishmarev, D., and P. W. Kuchel. 2016. NMR magnetization-transfer analysis of rapid membrane transport in human erythrocytes. Biophys. Rev. 8:369–384.

35. Montel-Hagen, A., M. Sitbon, and N. Taylor. 2009. Erythroid glucose transporters. Curr. Opin. Hematol. 16:165–172.

36. Raftos, J. E., B. T. Bulliman, and P. W. Kuchel. 1990. Evaluation of an electrochemical model of erythrocyte pH buffering using 31P nuclear magnetic resonance data. J. Gen. Physiol. 95:1183–1204.

37. Leitch, J. M., and A. Carruthers. 2009. α-and β-Monosaccharide transport in human erythrocytes. Am. J. Physiol.: Cell Physiol. 296:C151–C161.

38. Stein, W. D., and T. Litman. 2015. Channels, Carriers, and Pumps: an Introduction to Membrane Transport. Academic Press, San Diego.

39. Pagès, G., D. Szekely, and P. W. Kuchel. 2008. Erythrocyte-shape evolution recorded with fast-measurement NMR diffusion–diffraction. J. Magn. Reson. Imaging 28:1409–1416.

40. Wong, P. 2011. The basis of echinocytosis of the erythrocyte by glucose depletion. Cell Biochem. Funct. 29:708–711.

41. Young, J. D., S. Y. Yao, C. E. Cass, and S. A. Baldwin. 2003. Equilibrative nucleoside transport proteins. In Red Cell Membrane Transport in Health and Disease. I. Bernhardt, and J. C. Ellory, editors. Springer-Verlag, Berlin, pp. 321–337.

42. Grimes, A. 1980. Human Red Cell Metabolism. Blackwell Scientific Publications, Oxford.

43. Himmelreich, U., K. N. Drew, A. S. Serianni, and P. W. Kuchel. 1998. 13C NMR studies of vitamin C transport and its redox cycling in human erythrocytes. Biochemistry 37:7578–7588.

44. Sage, J. M., and A. Carruthers. 2014. Human erythrocytes transport dehydroascorbic acid and sugars using the same transporter complex. Am. J. Physiol.: Cell Physiol. 306:C910–C917.

45. May, J. M. 1998. Ascorbate function and metabolism in the human erythrocyte. Front. Biosci. 3:d1–10.

46. Bubb, W. A., H. A. Berthon, and P. W. Kuchel. 1995. Tris buffer reactivity with low-molecular-weight aldehydes: NMR characterization of the reactions of glyceraldehyde-3-phosphate. Bioorg. Chem. 23:119–130.

47. London, R. E., and S. A. Gabel. 1995. Fluorine-19 NMR studies of glucosyl fluoride transport in human erythrocytes. Biophys. J. 69:1814–1818.

48. Stein, W. D., and W. R. Lieb. 1986. Transport and Diffusion Across Cell Membranes. Academic Press, Orlando.

49. Carruthers, A., and D. Melchior. 1985. Transport of α- and β-D-Glucose by the intact human red cell. Biochemistry 24:4244–4250.

50. Janoshazi, A., and A. Solomon. 1993. Initial steps of α- and β-D-glucose binding to intact red cell membrane. J. Membr. Biol. 132:167–178.

51. Ginsburg, H. 1978. Galactose transport in human erythrocytes. The transport mechanism is resolved into two simple asymmetric antiparallel carriers. Biochim. Biophys. Acta, Biomembr. 506:119–135.

52. Ginsburg, H., and S. Yeroushalmy. 1978. Effects of temperature on the transport of galactose in human erythrocytes. J. Physiol. 282:399–417.

53. Leitch, J. M., and A. Carruthers. 2007. ATP-dependent sugar transport complexity in human erythrocytes. Am. J. Physiol.: Cell Physiol. 292:C974–986.

54. Faust, R. G. 1960. Monosaccharide penetration into human red blood cells by an altered diffusion mechanism. J. Cell. Physiol. 56:103–121.

55. Miwa, I., H. Fujii, and J. Okuda. 1988. Asymmetric transport of D-glucose anomers across the human erythrocyte membrane. Biochem. Int. 16:111–117.

56. Barnett, J. E., G. D. Holman, R. A. Chalkley, and K. A. Munday. 1975. Evidence for two asymmetric conformational states in the human erythrocyte sugar-transport system. Biochem. J. 145:417–429.

57. Appleman, J. R., and G. E. Lienhard. 1989. Kinetics of the purified glucose transporter. Direct measurement of the rates of interconversion of transporter conformers. Biochemistry 28:8221–8227.

58. Kuchel, P. W., B. E. Chapman, and J. R. Potts. 1987. Glucose transport in human erythrocytes measured using 13C NMR spin transfer. FEBS Lett. 219:5–10.

59. Barnett, J., G. Holman, and K. Munday. 1973. Structural requirements for binding to the sugar-transport system of the human erythrocyte. Biochem. J. 131:211–221.

60. Percival, M. D., and S. G. Withers. 1992. Binding energy and catalysis: deoxyfluoro sugars as probes of hydrogen bonding in phosphoglucomutase. Biochemistry 31:498–505.

61. Kayano, T., H. Fukumoto, R. Eddy, Y. S. Fan, M. Byers, T. B. Shows, and G. Bell. 1988. Evidence for a family of human glucose transporter-like proteins. Sequence and gene localization of a protein expressed in fetal skeletal muscle and other tissues. J. Biol. Chem. 263:15245–15248.

62. Müller, K., C. Faeh, and F. Diederich. 2007. Fluorine in pharmaceuticals: looking beyond intuition. Science 317:1881–1886.

63. Ametamey, S. M., M. Honer, and P. A. Schubiger. 2008. Molecular Imaging with PET. Chem. Rev. 108:1501–1516.

64. Soueidan, O.-M., B. J. Trayner, T. N. Grant, J. R. Henderson, F. Wuest, F. G. West, and C. I. Cheeseman. 2015. New fluorinated fructose analogs as selective probes of the hexose transporter protein GLUT5. Org. Biomol. Chem. 13:6511–6521.

65. Ellory, J. C., and J. D. Young. 1982. Red Cell Membranes: A Methodological Approach. Academic Press, London.

66. Moore, W. J. 1972. Physical Chemistry. Longman, Harlow, Essex, UK.

67. King, E. L., and C. Altman. 1956. A schematic method of deriving the rate laws for enzyme-catalyzed reactions. J. Phys. Chem. 60:1375–1378.

68. Orsi, B. A. 1972. A simple method for the derivation of the steady-state rate equation for an enzyme mechanism. Biochim. Biophys. Acta 258:4–8.

69. Indge, K. J., and R. E. Childs. 1976. New method for deriving steady-state rate equations suitable for manual or computer use. Biochem. J. 155:567–570.

70. Kuchel, P. W., and G. B. Ralston. 1988. Schaum’s Outline of Theory and Problems of Biochemistry. McGraw-Hill, New York.

71. Cornish-Bowden, A. 1977. An automatic method for deriving steady-state rate equations. Biochem. J. 165:55–59.

72. Mulquiney, P. J., and P. W. Kuchel. 2003. Modelling Metabolism with Mathematica. CRC Press, Boca Raton.

73. Britton, H. G. 1964. Permeability of the human red cell to labelled glucose. J. Physiol. 170:1–20.

74. Pagès, G., M. Puckeridge, G. Liangfeng, Y. L. Tan, C. Jacob, M. Garland, and P. W. Kuchel. 2013. Transmembrane exchange of hyperpolarized 13C-urea in human erythrocytes: subminute timescale kinetic analysis. Biophys. J. 105:1956–1966.

## Supporting References

1. Shishmarev, D., and P. W. Kuchel. 2016. NMR magnetization-transfer analysis of rapid membrane transport in human erythrocytes. Biophys. Rev. 8:369–384.

2. Raftos, J. E., B. T. Bulliman, and P. W. Kuchel. 1990. Evaluation of an electrochemical model of erythrocyte pH buffering using ^31^P nuclear magnetic resonance data. J. Gen. Physiol. 95:1183–1204.

3. Potts, J. R., and P. W. Kuchel. 1992. Anomeric preference of fluoroglucose exchange across human red-cell membranes: ^19^F-NMR studies. Biochem. J. 281:753–759.

4. O’Connell, T. M., S. A. Gabel, and R. E. London. 1994. Anomeric dependence of fluorodeoxyglucose transport in human erythrocytes. Biochemistry 33:10985–10992.

5. Stein, W. D., and W. R. Lieb. 1986. Transport and Diffusion Across Cell Membranes. Academic Press, Orlando.

6. Ellory, J. C., and J. D. Young. 1982. Red Cell Membranes: A Methodological Approach. Academic Press, London.

7. Moore, W. J. 1972. Physical Chemistry. Longman, Harlow, Essex, UK.

8. King, E. L., and C. Altman. 1956. A schematic method of deriving the rate laws for enzyme-catalyzed reactions. J. Phys. Chem. 60:1375–1378.

9. Orsi, B. A. 1972. A simple method for the derivation of the steady-state rate equation for an enzyme mechanism. Biochim. Biophys. Acta 258:4–8.

10. Indge, K. J., and R. E. Childs. 1976. New method for deriving steady-state rate equations suitable for manual or computer use. Biochem. J. 155:567–570.

11. Kuchel, P. W., and G. B. Ralston. 1988. Schaum’s Outline of Theory and Problems of Biochemistry. McGraw-Hill, New York.

12. Cornish-Bowden, A. 1977. An automatic method for deriving steady-state rate equations. Biochem. J. 165:55–59.

13. Mulquiney, P. J., and P. W. Kuchel. 2003. Modelling Metabolism with *Mathematica*. CRC Press, Boca Raton.

14. Britton, H. G. 1964. Permeability of the human red cell to labelled glucose. J. Physiol. 170:1–20.

15. Leitch, J. M., and A. Carruthers. 2009. α-and β-Monosaccharide transport in human erythrocytes. Am. J. Physiol.: Cell Physiol. 296:C151–C161.

16. Nelson, D. L., and M. M. Cox. 2008. Lehninger Principles of Biochemistry. W.H. Freeman and Company, New York.

17. Pagès, G., M. Puckeridge, G. Liangfeng, Y. L. Tan, C. Jacob, M. Garland, and P. W. Kuchel. 2013. Transmembrane exchange of hyperpolarized ^13^C-urea in human erythrocytes: subminute timescale kinetic analysis. Biophys. J. 105:1956–1966.

18. Ginsburg, H. 1978. Galactose transport in human erythrocytes. The transport mechanism is resolved into two simple asymmetric antiparallel carriers. Biochim. Biophys. Acta, Biomembr. 506:119–135.

19. Stein, W. D., and T. Litman. 2015. Channels, Carriers, and Pumps: an Introduction to Membrane Transport. Academic Press, San Diego.

20. London, R. E., and S. A. Gabel. 1995. Fluorine-19 NMR studies of glucosyl fluoride transport in human erythrocytes. Biophys. J. 69:1814–1818.

